# A genetically encoded red fluorescence dopamine biosensor enables dual imaging of dopamine and norepinephrine

**DOI:** 10.1101/2020.05.25.115162

**Authors:** Chihiro Nakamoto, Yuhei Goto, Yoko Tomizawa, Yuko Fukata, Masaki Fukata, Kasper Harpsøe, David E. Gloriam, Kazuhiro Aoki, Tomonori Takeuchi

## Abstract

Dopamine (DA) and norepinephrine (NE) are pivotal neuromodulators that regulate a broad range of brain functions, often in concert. Despite their physiological importance, untangling the relationship between DA and NE in finely controlling output functions is currently challenging, primarily due to a lack of techniques to visualize spatiotemporal dynamics with sufficiently high selectivity. Although genetically encoded fluorescent biosensors have been developed to detect DA, their poor selectivity prevents distinguishing DA from NE. Here, we report the development of a red fluorescent genetically encoded GPCR (G protein-coupled receptor)-activation reporter for DA termed ‘R-GenGAR-DA’. More specifically, a circular permutated red fluorescent protein (cpmApple) was inserted into the third intracellular loop of human DA receptor D1 (DRD1) followed by the screening of mutants within the linkers between DRD1 and cpmApple. We developed two variants: R-GenGAR-DA1.1, which brightened following DA stimulation, and R-GenGAR-DA1.2, which dimmed. R-GenGAR-DA1.2 demonstrated reasonable dynamic range (ΔF/F_0_ = –50%) and DA affinity (EC_50_ = 0.7 µM) as well as the highest selectivity for DA over NE (143-fold) amongst available DA biosensors. Due to its high selectivity, R-GenGAR-DA1.2 allowed dual-color fluorescence live imaging for monitoring DA and NE, combined with the existing green-NE biosensor GRABNE1m, which has high selectivity for NE over DA (>350-fold) in HeLa cells and hippocampal neurons grown from primary culture. By enabling precise measurement of DA, as well as simultaneous visualization of DA and NE, the red-DA biosensor R-GenGAR-DA1.2 is promising in advancing our understanding of the interplay between DA and NE in organizing key brain functions.

**Significance Statement:** The neuromodulators dopamine and norepinephrine modulate a broad range of brain functions, often in concert. One current challenge is to measure dopamine and norepinephrine dynamics simultaneously with high spatial and temporal resolution. We therefore developed a red-dopamine biosensor that has 143-fold higher selectivity for dopamine over norepinephrine. Taking advantage of its high selectivity for dopamine over norepinephrine, this red-dopamine biosensor allowed dual-color fluorescence live imaging for monitoring dopamine and norepinephrine in both HeLa cells and hippocampal neurons *in vitro* combined with the existing green-norepinephrine biosensor that has 350-fold selectivity for norepinephrine over dopamine. Thus, this approach can provide new opportunities to advance our understanding of high spatial and temporal dynamics of dopamine and norepinephrine in normal and abnormal brain functions.

## Introduction

The catecholaminergic neuromodulators dopamine (DA) and norepinephrine (NE) have very high structural similarity, differing only by a single hydroxy group. Dopaminergic projections mainly originate from the ventral tegmental area and substantia nigra pars compacta (1), whilst noradrenergic projections originate from the locus coeruleus (LC) (2, 3). It was discovered recently that noradrenergic LC axons co-released DA along with NE (4–6). DA is involved in reward (7, 8), motivation (9), novelty response (10), and motor control (11, 12). In addition, the involvement of DA and NA in many brain functions overlaps (13, 14), such as learning and memory (10, 15), arousal (16, 17), and stress response (6, 18). In particular, the prefrontal cortex receives both dopaminergic and noradrenergic projections, and these systems are involved in attention (19, 20) and working memory (21–23). Furthermore, dysfunction of dopaminergic or noradrenergic systems are thought to be associated with psychiatric disorders and neurodegenerative diseases, such as attention-deficit/hyperactivity disorder (ADHD), schizophrenia, and Parkinson’s disease (24–26).

Although interactions between dopamine and norepinephrine theoretically depend on the timing of release, spatial diffusions, concentrations, and neuromodulator ratios, little is actually known about these properties with high spatial and temporal resolution within the same preparation due to the technical limitations. For example, microdialysis with high-performance liquid chromatography has high sensitivity and selectivity to detect either DA and NE, but suffers from poor spatial and temporal resolution (27, 28). In contrast, fast-scan cyclic voltammetry (29) and a synthetic catecholamine nanosensor (30) have higher sensitivity and temporal resolution, but cannot distinguish between DA and NE. A method combining sensitivity, specificity, and spatiotemporal resolution is required to satisfactorily answer research questions regarding timing of release, spatial diffusions, concentrations, and ratios.

Recently developed genetically encoded fluorescent biosensors are able to detect extracellular DA or NE with high spatial and temporal resolution and sensitivity in freely moving animals using *in vivo* imaging (31–34). Binding of DA or NE to the sensor induces a conformational change, which couples with a change in the fluorescence of circular-permutated fluorescent protein, such as green fluorescent protein (GFP) for green fluorescence (31–34) and mApple for red fluorescence (34). The green-NE biosensor, GRAB_NE1m_ (abbreviated NE1m), has a high selectivity for NE (> 350-fold selectivity for NE over DA) (33). However, current DA biosensors do not have high enough selectivity for DA over NE [Green-DA biosensors: dLight1.1 (60-fold, but see *SI Appendix*, Fig. S7), GRAB_DA1h_ (∼ 10-fold), and GRAB_DA2m_ (15-fold); Red-DA biosensor: rGRAB_DA1m_ (22-fold)] (31, 32, 34) and consequently, it is difficult to use these DA biosensors for the simultaneous detection of DA and NE.

To image DA and NE dynamics simultaneously with high spatial and temporal resolution, we developed a red-DA biosensor using circular-permutated mApple (cpmApple), which has high selectivity for DA (143-fold selectivity for DA over NE). Using this red-DA biosensor with the existing green-NE biosensor, NE1m, which has high selectivity for NE (33), allowed us to successfully perform dual-color fluorescence monitoring of DA and NE with live imaging in both HeLa cells and primary culture of rat hippocampal neurons *in vitro*.

## Results

### Development and characterization of a red-DA biosensor

To develop a genetically encoded red fluorescence DA biosensor, we adopted the same approach as used to develop the green-DA sensors dLight (31) and GRAB_DA_ (32). First, we constructed an initial red-DA biosensor variant by inserting a red fluorescent protein, cpmApple (35), with linker sequences between Lys 232 and Lys 269 of human DA receptor D1 (DRD1), similarly to that done to construct dLight. We named it <u>r</u>ed fluorescent <u>g</u>enetically <u>en</u>coded <u>G</u>PCR <u>a</u>ctivation <u>r</u>eporter for DA, ‘R-GenGAR-DA1.0’ (abbreviated DA1.0; Fig. 1*A*). However, when DA was applied, DA1.0 did not exhibit a fluorescence response (*SI Appendix*, Fig. S1*A*). To improve its fluorescence response to DA, random mutagenesis was performed on the linker peptide sequences between DRD1 and cpmApple on DA1.0 (Fig. 1*A*). HeLa cells expressing mutants of DA1.0 were stimulated by application of DA, and the change in red fluorescence intensity was quantified (Fig. 1*B*). Of 864 mutants, we selected three mutants (#76, #310, and #430) that responded positively to DA and subjected them to time-lapse imaging (Fig. 1 *C* and *D*, and *SI Appendix*, Fig. S1*B*). All three mutants showed detectable red fluorescence increases in response to DA application and this response was blocked by the DRD1/5 antagonist SCH 23390 (SCH) (Fig. 1 *C* and *D*, and *SI Appendix*, Fig. S1*B*). The amino acid sequences of mutated linkers were determined in these mutants (*SI Appendix*, Fig. S1*C*). We selected #76 (‘R-GenGAR-DA1.1’, abbreviated DA1.1) because it showed the largest positive response to DA amongst the three mutants. We then characterized the dose-response curves of DA1.1 for DA and NE and calculated the half maximal effective concentration (EC_50_). As a result, DA1.1 showed 12.6-fold selectivity for DA over NE (Fig. 1*E*).

**Fig. 1.**
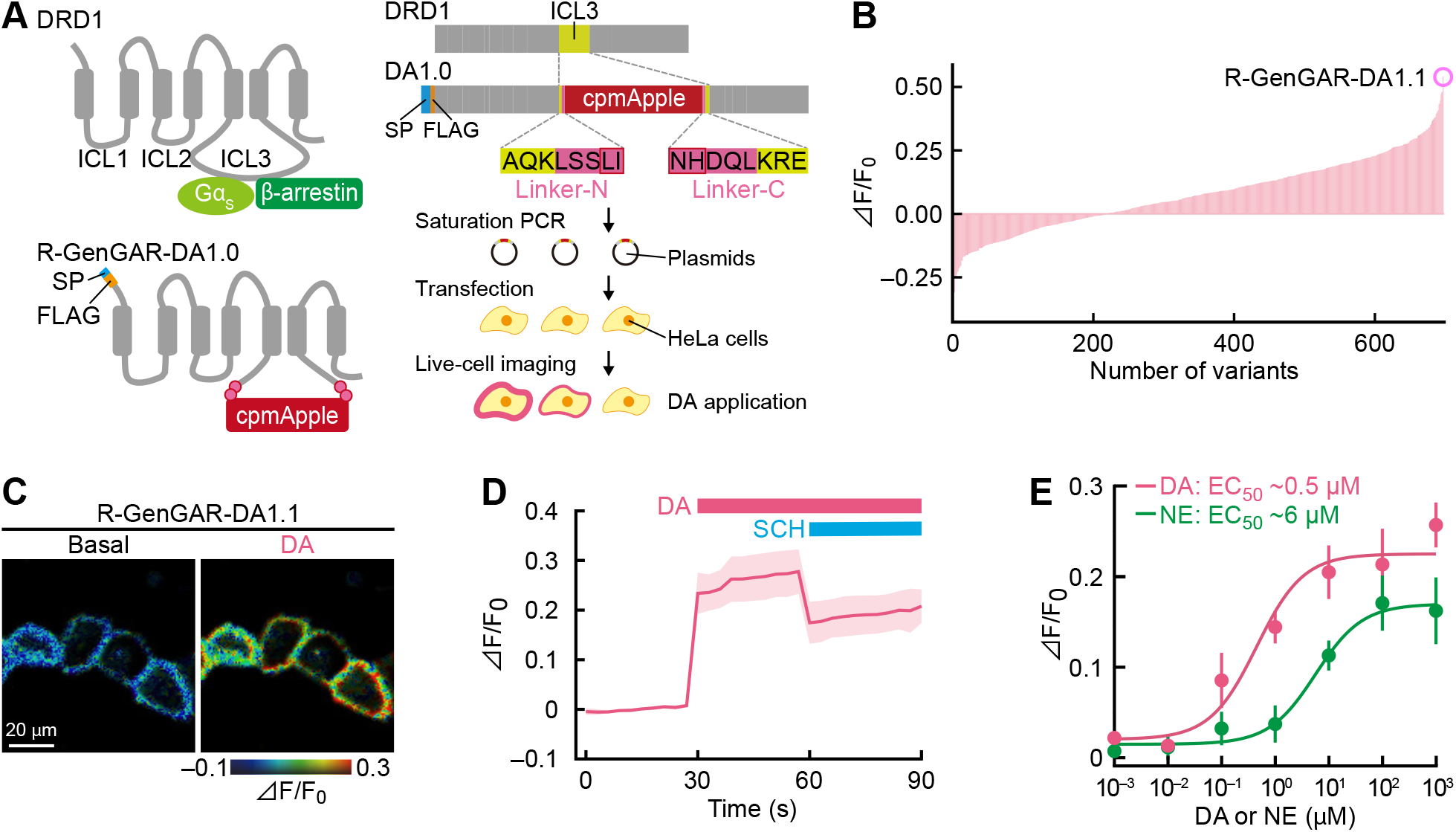
Development of R-GenGAR-DA1.1, which showed a positive response to dopamine (DA). (*A*) Strategy to develop R-GenGAR-DA1.0. Left panels show a schematic illustration of human DRD1 and the red fluorescent protein ‘cpmApple’ insertion site. Right panels show screening flow chart. Linker sequences connecting DRD1 and cpmApple were randomly mutated using saturation PCR. The plasmids expressing each linker mutant were isolated, followed by the transfection into HeLa cells by lipofection. Changes in fluorescence intensity following 10 μM DA stimulation was monitored by live-cell imaging of HeLa cells expressing each mutant. ICL, intracellular loop; SP, signal peptide (hemagglutinin secretary sequence). (*B*) Summary of screening results. The normalized fluorescence changes (ΔF/F_0_) of the HeLa cells expressing each mutant in response to 10 μM DA stimulation are shown. Each bar represents the average of 1-3 independent experiments. We selected a mutant “R-GenGAR-DA1.1” that showed a maximum response to 10 μM DA stimulation. (*C*) Representative images of HeLa cells expressing DA1.1 stimulated with 10 µM DA. The fluorescence change (ΔF/F_0_) before and after DA stimulation are shown in the pseudocolor intensity-modulated display mode. (*D*) Normalized fluorescence change (ΔF/F_0_) of DA1.1 in HeLa cells in panel *C*. DA (10 µM) and SCH 23390 (SCH, 10 µM) were treated at the time points indicated by pink and blue bars, respectively (*SI Appendix*, Fig. S5*A*). Mean ΔF/F_0_ values of 10 cells from 1 experiment are shown with SD (shaded area). (*E*) Dose-response curves, with temperature equilibration, of DA (pink) and NE (green) in HeLa cells expressing DA1.1 (*SI Appendix*, Fig. S4*D*). DA: max ΔF/F_0_ = 0.23 ± 0.02% and EC_50_ = 0.45 ± 0.21 µM; NE: max ΔF/F_0_ = 0.17 ± 0.03% and EC_50_ = 5.68 ± 1.19 µM (DA and NE, *n* = 3 experiments in both cases, 10 cells per experiment). Experimental data (dots) were fitted with the Hill equation (lines). DA1.1 has 12.6-fold selectivity for DA over NE.

### Development and characterization of an inverse-type red-DA biosensor

The dynamic range of DA1.1 and its selectivity for DA were lower than some other DA biosensors (31, 32, 34), prompting us to make further improvements. We attempted to expand the dynamic range of DA1.1 by introducing the same mutations as in the green-DA biosensor dLight1 (31). Substitution of Phe 129 with Ala (F129A mutation of dLight1.2; *SI Appendix*, Fig. S2*A*), and addition of Glu to the N-terminal linker (dLight1.3a; Fig. 2*A*) were previously shown to significantly increase the dynamic range of dLight1.1 (31). The F129A mutation in DA1.1, however, led to only a slight increase in the fluorescent signal upon DA application (*SI Appendix*, Fig. S2*B*) and its sensitivity to both DA and NE was lower than that of the original DA1.1 (data not shown). Surprisingly, the addition of Glu to the N-terminal linker in DA1.1 showed bright red fluorescence in the basal state. This variant had substantially reduced fluorescence signal in response to DA, which subsequently returned to basal level following treatment with SCH (Fig. 2*B*). We named this inverse type red fluorescence DA biosensor ‘R-GenGAR-DA1.2’ (abbreviated DA1.2).

**Fig. 2.**
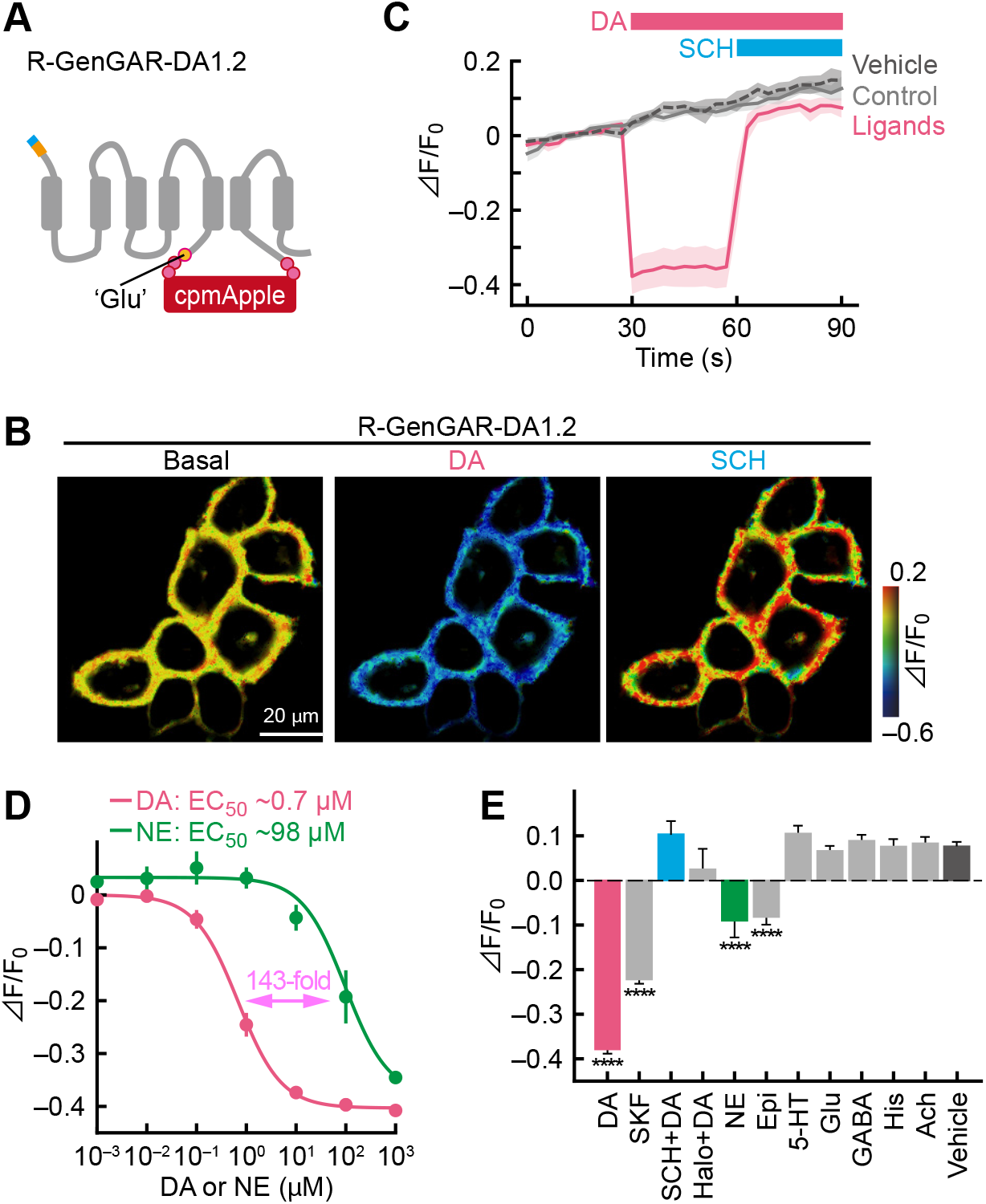
Development of R-GenGAR-DA1.2, which showed a negative red fluorescence response to DA. (*A*) Schematic illustration of a mutation site. Glutamate (Glu) was introduced into the N-terminal side of a linker in DA1.1. (*B*) Representative images of HeLa cells expressing DA1.2 treated with 10 µM DA, followed by 10 µM DRD1/5 antagonist SCH 23390 (SCH). Images are shown in the pseudocolor intensity-modulated display mode. (*C*) Normalized fluorescence change (ΔF/F_0_) of DA1.2 in HeLa cells. Medium temperature was equilibrated before imaging (*SI Appendix*, Fig. S5*B*). DA (10 µM) and SCH (10 µM) were treated at the time points indicated by pink and blue bars, respectively. Mean ΔF/F_0_ of 30 cells from 3 experiment are shown with SD (shaded area). Vehicle, 10 µM HCl or water; control, cells were only exposed to emission light. (*D*) The dose-response curves, with temperature equilibration, of DA (pink) and NE (green) on HeLa cells expressing DA1.2 (*SI Appendix*, Fig. S4*D*). DA: max ΔF/F_0_ = –0.40 ± 0.01% and EC_50_ = 0.68 ± 0.08 µM; NE: max ΔF/F_0_ = –0.45 ± 0.05% and EC_50_ = 98 ± 51 µM (DA and NE, *n* = 4 experiments in both cases). Experimental data (dots) were fitted with the Hill equation (lines). DA1.2 has 143-fold selectivity for DA over NE. (*E*) Selectivity of DA1.2 for pharmacological compunds (*n* = 3-4 experiments, 10 cells per experiment; *SI Appendix*, Fig. S5*C*). All compounds were 10 µM. DRD1 agonist SKF 81297 (SKF), SCH, DRD2 antagonist haloperidol (Halo), epinephrine (Epi), serotonin (5-HT), glutamate (Glu), γ-aminobutyric acid (GABA), histamine (His), and acetylcholine (ACh). For the vehicle condition, there was no significant difference between 10 µM HCl in H_2_O and 0.001% dimethyl sulphoxide (DMSO) (*n* = 4 experiments in each, 10 cells per experiment; Mann-Whitney *U*-test, *P* = 0.69). Therefore, these values were averaged and used as the vehicle condition. Mean ΔF/F_0_ values are shown with SEM. One-way ANOVA, *F*_12,38_ = 53.11, *P* < 0.0001; Dunnett’s post hoc test (vs vehicle), \*\*\*\**P* < 0.0001.

Unexpectedly, time-lapse imaging of DA1.2 in HeLa cells showed that the baseline fluorescent signals increased gradually in both vehicle and control conditions (data not shown). We explored the cause of this phenomenon and found that the DA1.2 baseline fluorescent signals were seemingly associated with thermochromism and photochromism stemming from cpmApple (36) (*SI Appendix*, Fig. S3). The former effect, thermochromism, was apparent from the inverse relationship between baseline fluorescent signals of R-GenGAR-DA and incubation temperature (*SI Appendix*, Fig. S3 *A–D*). Thermochromism was observed when test compounds were added to our experimental system. Therefore, imaging was performed after temperature equilibration (*SI Appendix*, Fig. S4 and Fig. S5). With respect to photochromism, irradiating light at wavelengths of either 488 nm or 561 nm induced an increase in basal red fluorescence of DA1.2 expressed in HeLa cells under constant medium temperature (*SI Appendix*, Fig. S3*E*). This effect was most pronounced when the irradiation light was at full power. Because light of 488 nm and 561 nm is generally used to excite green and red fluorophores respectively, it was problematic that the photochromism on DA1.2 was induced by the irradiation of those light wavelengths, especially when we combined DA1.2 with another fluorescent biosensor to perform dual-color time-lapse imaging in HeLa cells and primary hippocampal neurons (Fig. 3 and Fig. 5). We found that the increased baseline fluorescence observed in DA1.2 in primary hippocampal neurons was reduced following irradiation of two streams of light (561 nm followed by 488 nm) for 150 s (*SI Appendix*, Fig. S3*F*). Therefore, pre-light exposure just prior to dual-color time-lapse imaging was conducted to reduce the photochromism effect in DA1.2 (Fig. 3 and Fig. 5, *SI Appendix*, Fig. S5*D* and Table S1). Time-lapse imaging of DA1.2 with temperature equilibration (*SI Appendix*, Fig. S5*B*) showed that DA application lowered red fluorescence, which was restored following SCH treatment (Fig. 2*C*). Baseline fluorescence intensity still increased moderately in both vehicle and control conditions, possibly due to photochromism. The dose-response curve with temperature equilibration (*SI Appendix*, Fig. S4*D–F*) shows that DA1.2 has a slightly higher dynamic range and comparable affinity to DA (max ΔF/F_0_ = – 0.40 ± 0.01% and EC_50_ = 0.68 ± 0.08 μM) compared to that of DA1.1 (Fig. 2*D*). It is of note that the selectivity of DA1.2 for DA over NE was 143-fold, much higher than that of DA1.1 (Fig. 2*D*), which was due to the decrease in the affinity of DA1.2 to NE. In order to further enhance the selectivity of DA1.2 for DA over NE, we attempted to predict mutations based on ligand-receptor structure models (see *SI Appendix*, Materials and Methods). Since the difference between DA and NE is only one additional hydroxy group on NE, preference for DA might be accomplished by making the area around the binding site unfavorable for this hydroxy group. Based on a structural model complex of the DRD1 with either DA or NE in the binding site (*SI Appendix*, Fig. S6*A*), we then introduced 13 mutations to DA1.2 designed to increase preference for DA over NE (*SI Appendix*, Fig. S6*B*). Six of the 13 mutants showed a change in red fluorescence response to DA (*SI Appendix*, Fig. S6 *C* and *D*). Although the dose-response curves with temperature equilibration for both DA and NE were obtained from three mutants, none of these demonstrated an increase in selectivity for DA over NE compared to that of DA1.2 (*SI Appendix*, Fig. S6*E*). We then confirmed that the selectivity of the red-DA biosensor DA1.2 (143-fold) was higher than that of green-DA biosensors dLight1.1 (17-fold), dLight1.2 (32-fold), and dLight1.3a (19-fold) in our experimental conditions (*SI Appendix*, Fig. S7). Consequently, 143-fold selectivity for DA over NE in DA1.2 is the highest amongst currently available DA biosensors (31, 34).

**Fig. 3.**
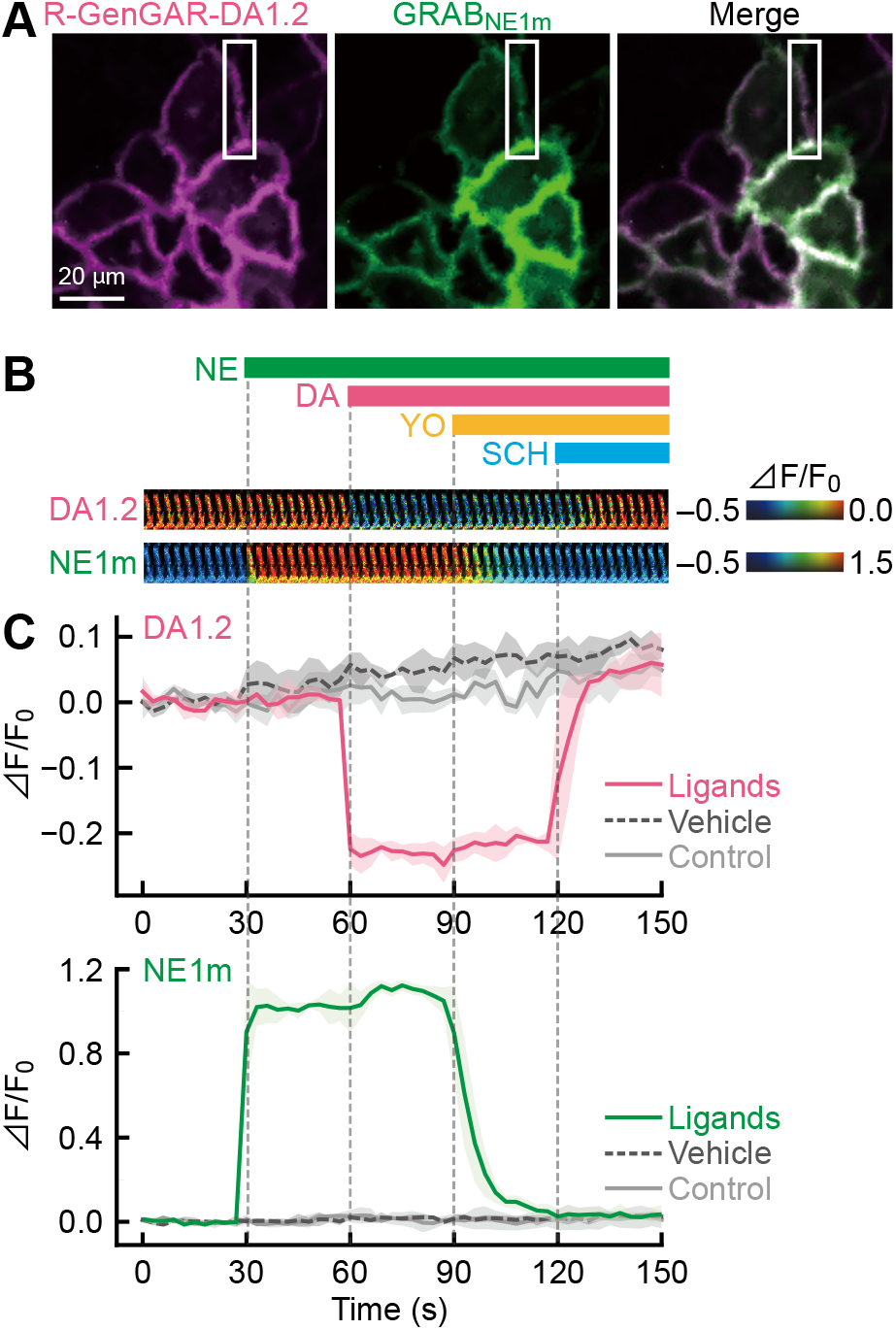
Dual-color fluorescence time-lapse imaging of R-GenGAR-DA1.2 combined with GRAB_NE1m_ in HeLa cells. (*A*) Representative image of HeLa cells co-expressing DA1.2 and NE1m. (*B*) Enlarged time-lapse images in the pseudocolor intensity-modulated display mode from the white boxed regions shown in panel *A*. Bars show the schedule of agonist/antagonist application to both DA1.2 and NE1m. Gray vertical lines indicate time of application. Concentrations: DA and SCH, 5 µM; NE and YO, 1 µM. Medium temperature and photochromism were equilibrated before imaging (*SI Appendix*, Fig. S5*D*). (*C*) Normalized fluorescence intensity change (ΔF/F_0_) of DA1.2 (top) and NE1m (bottom) in HeLa cells co-expressing DA1.2 and NE1m. Vehicle, 10 µM HCl in H_2_O or 0.001% DMSO; control, cells were only exposed to emission light. Mean ΔF/F_0_ values of 30 cells from 3 experiments are shown with SD (shaded areas). Result of statistical test is shown in *SI Appendix*, Fig. S9 *A* and *B*.

**Fig. 5.**
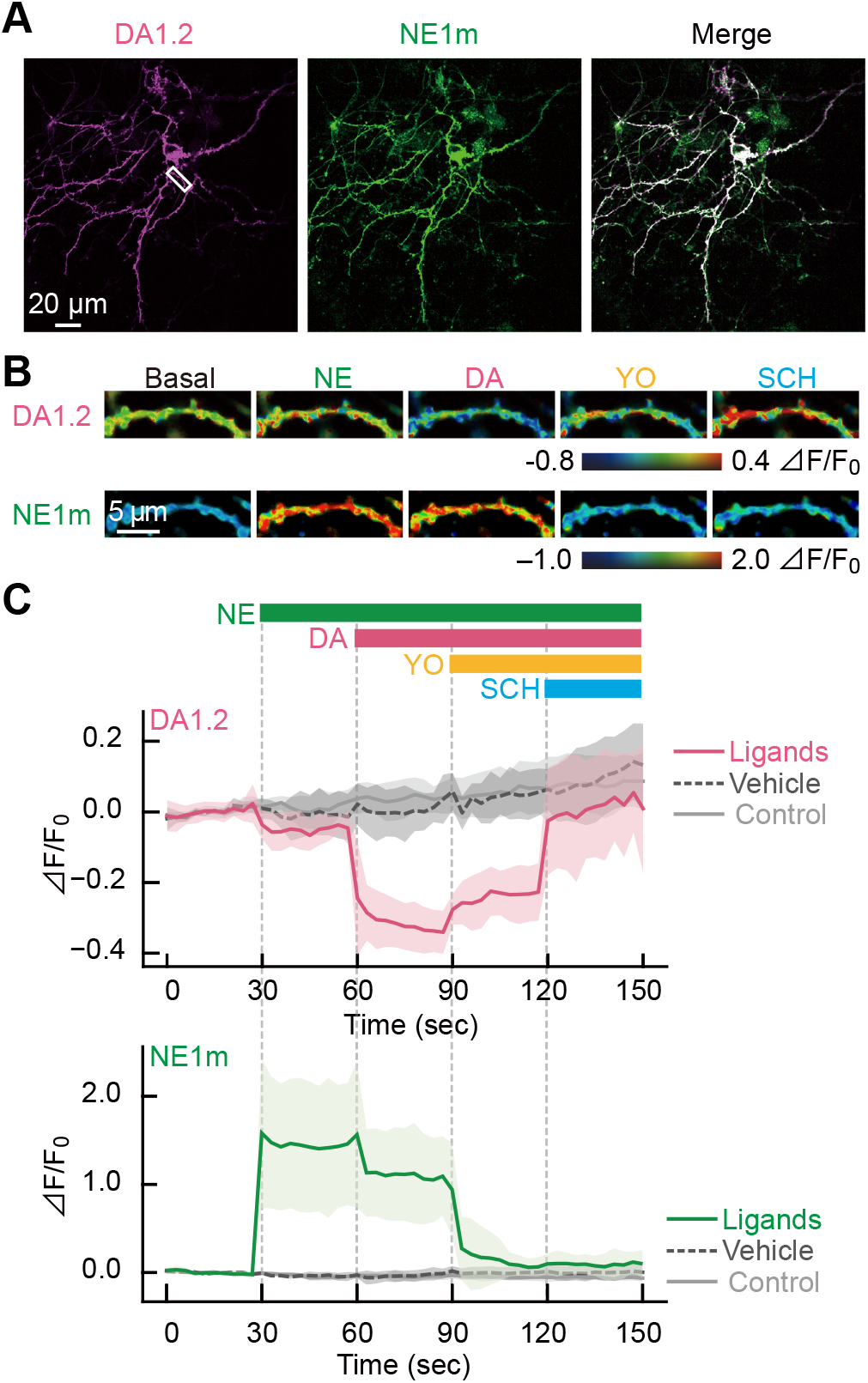
R-GenGAR-DA1.2 combined with GRAB_NE1m_ enables dual-color fluorescence imaging of DA and NE in a primary culture of rat hippocampal neurons. (*A*) Representative image of a primary hippocampal neuron co-expressing DA1.2 and NE1m. (*B*) Enlarged time-lapse images of DA1.2 and NE1m treated with agonists or antagonists in pseudocolor intensity-modulated display mode from the dendritic region in the primary hippocampal neurons marked as the white boxes in panel *A*. Concentrations: DA and SCH, 5 µM; NE and YO, 1 µM. Medium temperature and photochromism were equilibrated before imaging (*SI Appendix*, Fig. S5*D*). (*C*) Normalized fluorescence intensity change (ΔF/F_0_) of DA1.2 (top) and NE1m (bottom) in the primary hippocampal neurons co-expressing DA1.2 and NE1m. Bars show the schedule of agonist/antagonist application to both DA1.2 and NE1m. Gray vertical lines indicate time of application. Vehicle, 10 µM HCl in H_2_O or 0.001% DMSO; control, cells were only exposed to emission light. Colored lines indicate mean ΔF/F_0_ and light-colored shaded area is the SD. Ligands, 6 neurons from experiments; vehicles, 4 neurons from 4 experiments; control, 4 neurons from 1 experiment. Statistical test results are shown in *SI Appendix*, Fig. S9 *C* and *D*.

We further characterized how DA1.2 responded to a variety of test compounds. The DRD1/5 agonist SKF 81297 led to a partial response from DA1.2, whilst the response to DA was blocked by the DRD1/5 antagonist SCH (Fig. 2*E*). The application of several other neurotransmitters/neuromodulators showed no significant response in DA1.2 (Fig. 2*E*). In addition, we confirmed that DA1.2 induced no cyclic adenosine monophosphate (cAMP) increase upon DA application, indicating that, unlike wild-type DRD1, DA1.2 activity does not activate the canonical Gαs signaling pathway (*SI Appendix*, Fig. S8).

### Dual-color fluorescence imaging of DA and NE in HeLa cells

We then tested the simultaneous imaging of DA and NE at the single-cell level. To accomplish this, both DA1.2 and a green-NE sensor, NE1m, which has high selectivity for NE over DA (> 350-fold) (33), were co-expressed in HeLa cells (Fig. 3*A*). Following irradiation of DA1.2 by two streams of light (561 nm followed by 488 nm) to reduce the effects due to photochromism, we applied the following compounds in this order: NE (1 µM), DA (5 µM) followed by the α-adrenoceptor antagonist yohimbine (YO, 1 µM), and SCH (5 µM) (Fig. 3*B* and *SI Appendix*, Fig. S5*D*). As we expected, DA1.2 exhibited a decrease in fluorescence to DA, but not to NE, and its response to DA was blocked by SCH treatment (Fig. 3 *B* and *C* and *SI Appendix*, Fig. S9 *A* and *B*), confirming that the decrease in fluorescence could indeed be attributed to DA binding to DA1.2. Meanwhile, NE1m showed an increase in green fluorescence upon application of NE, but not DA, and its fluorescence response recovered to its basal level following YO treatment (Fig. 3 *B* and *C* and *SI Appendix*, Fig. S9 *A* and *B*). In summary, we demonstrated that our red-DA biosensor DA1.2 and the existing green-NE biosensor NE1m can distinguish DA and NE, respectively, in HeLa cells.

### Dual-color fluorescence imaging of DA and NE in a primary culture of rat hippocampal neurons

To further test the application of DA1.2, we introduced DA1.2 into rat primary hippocampal neurons, where it was successfully expressed, and distributed in plasma membranes throughout neurons (Fig. 4*A*). Application of DA (5 µM) led to reduced DA1.2 red fluorescence, and this was restored to baseline by SCH treatment (5 µM) (Fig. 4 *A* and *B*). Conversely, pretreatment with SCH completely suppressed the response of DA1.2 to DA (Fig. 4*C*), indicating that DA1.2 was successful in facilitating the visualization of DA in primary hippocampal neurons. The dose-response curve for DA with temperature equilibration was obtained in primary hippocampal neurons expressing DA1.2. In this set up, DA1.2 showed max ΔF/F_0_ = –0.51 ± 0.05%, and an EC_50_ value of 0.56 ± 0.01 μM (Fig. 4*D*), which were comparable to the results in HeLa cells.

**Fig. 4.**
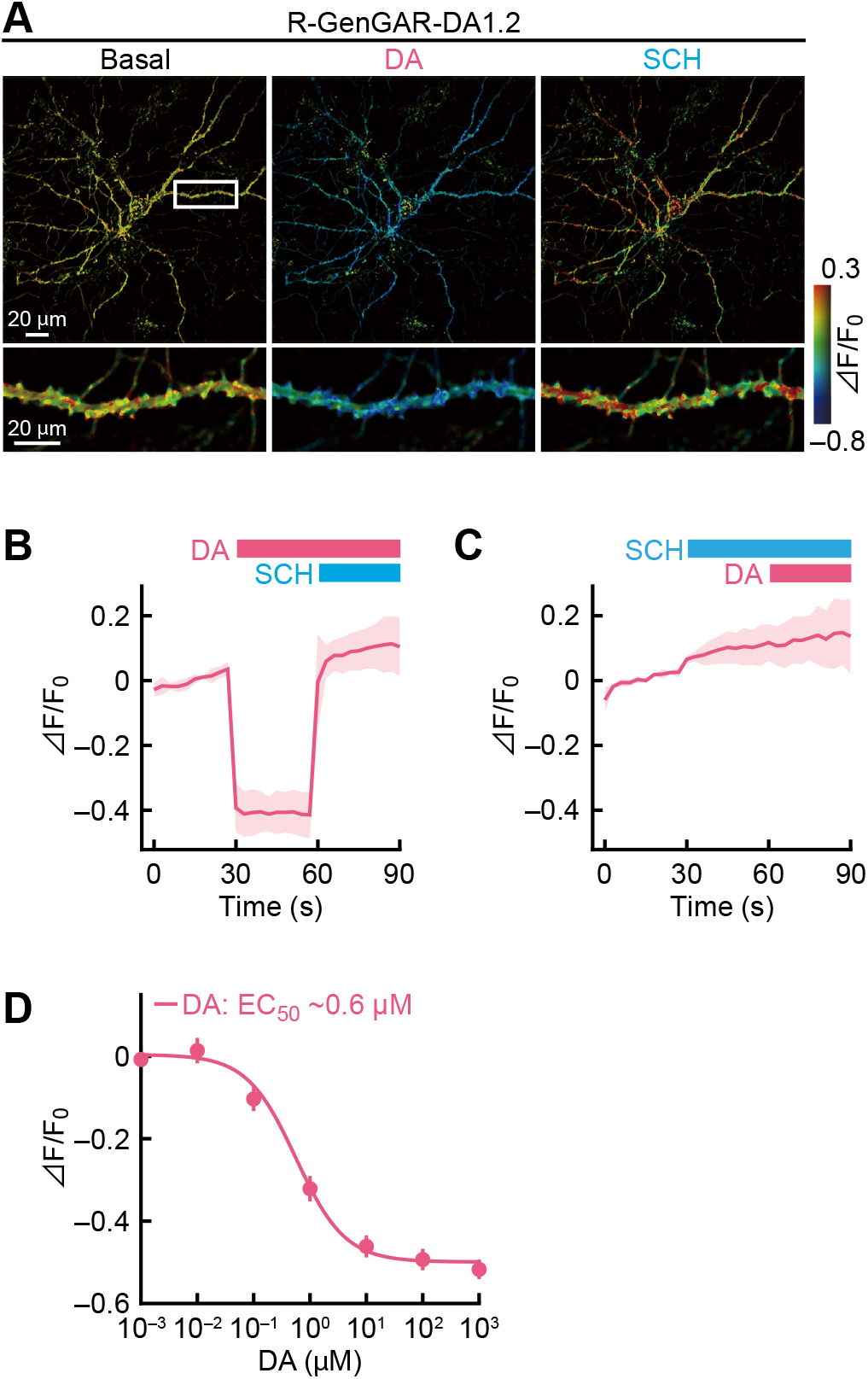
Characterization of R-GenGAR-DA1.2 in the primary culture of rat hippocampal neurons. (*A*) Representative images of a primary hippocampal neuron expressing DA1.2. The fluorescence change (ΔF/F_0_) before (left) and after the application of 5 μM DA (middle) followed by 5 μM SCH (right) (*SI Appendix*, Fig. S5*B*) are shown in pseudocolor intensity-modulated display mode. Bottom: magnification of dendrite marked in the top left image (white rectangle). Medium temperature was equilibrated before imaging. (*B*) Normalized fluorescence change (ΔF/F_0_) of DA1.2 in the primary hippocampal neurons in panel *A*. DA (5 μM) and SCH (5 μM) were treated at the time points indicated by pink and blue bars, respectively. Mean ΔF/F_0_ values of 6 neurons from 6 experiments are shown with SD (shaded area). (*C*) DA1.2 was pre-treated with SCH before application of DA. Mean ΔF/F_0_ of 3 neurons from 3 experiments are shown with SD (shaded area). Medium temperature was equilibrated before imaging. (*D*) The dose-response curve with temperature equilibration of DA (pink) on the primary hippocampal neurons expressing DA1.2 (*SI Appendix*, Fig. S4*D*). DA: max ΔF/F_0_ = –0.51 ± 0.05% and EC_50_ = 0.56 ± 0.01 µM; *n* = 7 neurons from 7 experiments. Experimental data (dots) were fitted with the Hill equation (lines).

We finally performed dual-color fluorescence imaging of DA and NE in the primary culture of rat hippocampal neurons. DA1.2 and NE1m were co-expressed in the primary hippocampal neurons (Fig. 5*A*). After the effects of photochromism were reduced using two streams of light (561 nm followed by 488 nm), we then applied compounds in the following order: NE (1 µM), DA (5 µM) followed by YO (1 µM), and SCH (5 µM) (*SI Appendix*, Fig. S5*D*). As we observed in HeLa cells, DA, but not NE application, led to a decrease in the red fluorescence signal of DA1.2, which was restored following SCH treatment (Fig. 5 *B* and *C* and *SI Appendix*, Fig. S9 *C* and *D*). In addition, we observed a NE-induced increase in NE1m green fluorescence, and this fluorescence response was blocked by YO treatment (Fig. 5 *B* and *C* and *SI Appendix*, Fig. S9 *C* and *D*).

## Discussion

We have developed genetically encoded red fluorescent DA sensors R-GenGAR-DA1.1 and DA1.2, which respond positively and negatively to DA, respectively. Specifically, DA1.2 demonstrated reasonable dynamic range (ΔF/F_0_ = –50%) and DA affinity (EC_50_ = 0.7 µM) as well as high selectivity for DA (143-fold higher affinity than for NE). In HeLa cells, dual-color live imaging of DA and NE was successfully performed using DA1.2 combined with the existing green-NE biosensor NE1m, which has high selectivity for NE over DA (> 350-fold) (33). Furthermore, DA1.2 and NE1m were also co-expressed in the primary culture of rat hippocampal neurons, allowing dual-color live imaging of DA and NE *in vitro*. We thus successfully demonstrated that application of two different color-based fluorescent neurochemical sensors (i.e. cpGFP- and cpmApple-based sensors) with high selectivity for each ligand allow us to monitor two different neurochemicals simultaneously.

A striking feature of DA1.2 is its high selectivity for DA over NE. For a DA biosensor, selectivity for DA over NE is critical to avoid cross-reactivity for imaging in the brain areas where NE is present at relatively higher amount than DA (5, 28, 37, 38). In our experimental conditions, the specificity for DA over NE, shown by green-DA biosensor dLight1 variants, was lower than that in a previous study (31): dLight1.1 (17-fold), dLight1.2 (32-fold), dLight1.3a (19-fold) (*SI Appendix*, Fig. S7). Sun and colleagues reported that DA biosensors GRAB_DA_ also did not have enough selectivity for DA over NE: GRAB_DA1h_ (∼10-fold), GRAB_DA2m_ (15-fold), and rGRAB_DA1m_ (22-fold) (32, 34). Thus, DA1.2 has the highest selectivity for DA over NE (143-fold) compared to all other currently available DA biosensors. Although we tried to further increase the selectivity of DA1.2 by introducing mutations predicted from *in silico* models (*SI Appendix*, Fig. S6), the selectivity of DA1.2 was not raised above the already obtained 143-fold level, possibly due to the use of models in the absence of the crystal structure of DRD1. Taking advantage of this high selectivity of DA1.2 for DA over NE, we succeeded in detecting DA and NE simultaneously in HeLa cells and primary hippocampal neurons *in vitro* by dual-color imaging combined with the existing green-NE biosensor NE1m, which has the highest selectivity for NE over DA (33). The affinity of DA1.2 to DA (EC_50_ = 0.68 µM in HeLa cells, and EC_50_ = 0.56 µM in primary hippocampal neurons) was at sub-micromolar levels, which is within the range of available DA biosensors, comparable to the dLight1 series (31) and lower than the GRAB_DA_ series (32, 34). In addition, other advantages of the red fluorescent DA1.2 sensor are lower phototoxicity and higher tissue penetration because of its longer excitation wavelength. Furthermore, DA1.2 enables multiplex imaging with other colored biosensors for different neurochemicals (39, 40), optogenetic actuators (41), intracellular signaling biosensors (42, 43), calcium indicators (44, 45), and voltage indicators (46).

Despite the successful simultaneous imaging of DA and NE using DA1.2 combined with NE1m *in vitro*, further improvements to DA1.2 will be required for use in *in vivo* imaging. The main areas in which improvement are required are: [i] expanding the dynamic range and [ii] lowering thermochromism and photochromism. The first concern is the relatively low dynamic range DA1.2 (ΔF/F_0_ ∼ –50%). Recent literature on biosensor development has shown that an increase in dynamic range can be achieved by optimization of the linker insertion site, linker length, and, by random mutagenesis, the amino acid sequences on the linkers and the circular-permutated fluorescent protein (34, 47, 48). In order to expand the dynamic range of DA1.2, these strategies should be applied to DA1.2 in future work. The second concern regarding DA1.2 is thermochromism and photochromism, which is due to cpmApple (36). The thermochromic effect could potentially be avoided if the temperature of the animal is kept constant during *in vivo* imaging. In addition, it was recently reported that photochromism due to a cpmApple was successfully diminished by introduction of 22 mutations in the cpmApple region of red-dopamine sensor rGRAB_DA_ (34). Therefore, it may be possible to minimize photochromism in DA1.2 by introducing these mutations into cpmApple. Once those issues are overcome, an improved DA1.2 could be an extremely useful tool for simultaneous measurements of extracellular DA and NE dynamics in the brains of freely moving animals.

Recently, there has been an increased demand for the development of tools to observe DA and NE dynamics simultaneously with high spatial and temporal resolution *in vivo*. For example, it was reported that pharmacological blockade of dopamine D_1_/D_5_ receptors in the hippocampus prevented a memory-boosting effect induced by environmental novelty or by optogenetic activation of noradrenergic LC neurons in mice (4). Later, Kempadoo and colleagues directly detected co-release of DA along with NE after optogenetic stimulation of LC axons in the hippocampus *ex vivo* using high-performance liquid chromatography (5). These discoveries raise many questions regarding the co-release of DA and NE from LC terminals into the hippocampus in freely moving animals. For instance, what are the ranges of spatial diffusion and the precise time courses of concentration change? High spatial and temporal dual-color imaging of DA and NE dynamics in the hippocampus could give us an opportunity to answer these questions. Furthermore, when fiber photometry or two-photon microscopy is applied, the dual-color imaging will enable the measurement of DA and NE at the same spot in the brain. Because of this, the extracellular spatiotemporal dynamics of DA and NE will be comparable to each other under the same conditions.

To the best of our knowledge, this is the first time that simultaneous live imaging of extracellular DA and NE has been performed with dual-color fluorescence in both HeLa cells and in a primary culture of rat hippocampal neurons *in vitro*. Here, this was accomplished using our red-DA biosensor DA1.2 combined with the existing green-NE biosensor NE1m. This approach will be able to provide new insights into the high spatial and temporal dynamics of neuromodulators DA and NE in brain areas of interests, leading to advances in our understanding of the mechanisms of interplay between DA and NE in organizing key brain functions. A better understanding of these neuromodulatory systems would have the potential to facilitate new ways of treating psychiatric disorders and neurodegenerative diseases.

## Materials and Methods

Animal experiments were approved by the Animal Care Committee of the National Institutes of Natural Sciences in Japan (19A029) and were performed in accordance with its guidelines. Details on animals, and procedures regarding drugs, molecular cloning, saturation polymerase chain reaction (PCR) for the screening of optimal linker sequences, design of DRD1 mutations, cell culture, drug administrations, fluorescence imaging, detection of cAMP signaling using cAMP biosensor, quantification of imaging and data analysis, and statistical analysis are detailed in *SI Appendix*, Materials and Methods.

## ACKNOWLEDGMENTS

We thank all members of the Aoki and Takeuchi laboratories for helpful discussions and assistance. Sachiko Furukawa and Yuri Miyazaki helped with the primary culture of rat hippocampal neurons. The plasmid DNA of GRAB_NE1m_, RGECO1, and DRD1-Tango was provided by Yulong Li, Takeharu Nagai, and Bryan Roth, respectively. We also thank Takanari Inoue, Adrian Duszkiewicz, David Bett, Mai Iwasaki, Yulong Li, Steffen Sinning, and Lina Bukowski for scientific discussion.

This work was supported by Japan Society for the Promotion of Science (JSPS) KAKENHI Grants (no.19K16050 to Y.G.; no. 19H03331 to Y.F.; no. 18H04873 to M.F.); Lundbeckfonden (R163-2013-16327) and Danmarks Frie Forskningsfond | Natural Sciences (8021-00173B) (to D.E.G.); Core Research for Evolutional Science and Technology | Japan Society for the Promotion of Science (JPMJCR1654), JSPS KAKENHI Grants (no. 16KT0069, 16H01425 “Resonance Bio”, 18H04754 “Resonance Bio”, 18H02444, and 19H05798), and ONO Medical Research Foundation (to K.A.); Novo Nordisk Fonden Young Investigator Award 2017 (NNF17OC0026774), Aarhus Institute of Advanced Studies (AIAS)-EU FP7 Cofund programme (754513), Lundbeckfonden (DANDRITE-R248-2016-2518), and the Cooperative Study Program (20-105) of National Institute for Physiological Sciences (to T.T.).

## Supplementary Information

### Materials and Methods

#### Animals

Pregnant Wistar/ST rats were purchased from Japan SLC, Inc. for the primary cultures of rat hippocampal neurons. All experiments were approved by the Animal Care Committee of the National Institutes of Natural Sciences in Japan (19A029), and were performed in accordance with its guidelines.

#### Compounds used to test fluorescence response

Dopamine (DA) hydrochloride (1 M stock, H8602, Sigma-Aldrich), serotonin hydrochloride (50 mM stock, 14332, CAY,), and L-adrenaline (epinephrine) (5 mM stock, A0173, TCI) were dissolved in 10 mM HCl. L-noradrenaline bitartrate monohydrate (1 M stock, A0906, TCI), sodium L-glutamate monohydrate (10 mM stock, G0188, TCI), 4-aminobutyric acid (100 mM stock, A0282, TCI), histamine (100 mM stock, 18111-71, Nacalai Tesque), acetylcholine chloride (10 mM stock, A6625, Sigma-Aldrich), and R(+)-SCH 23390 hydrochloride (10 mM stock, D054, Sigma-Aldrich) were each dissolved separately in distilled water. SKF 81297 hydrobromide (10 mM stock, 1447, TOCRIS), haloperidol hydrochloride (20 mM stock, 0931, TOCRIS), and yohimbine hydrochloride (20 mM stock, 1127, TOCRIS) were dissolved in DMSO. Compound solutions were then subdivided into aliquots and stored at −20 °C until use. A working solution of 1 M DA was stored at 4 °C for 3 weeks prior to use.

#### Plasmids

R-GenGAR-DA1.0 cDNA and dLight1.1 (Patriarchi *et al*., 2018) cDNA were synthesized by FASMAC Co. Ltd. into the vector plasmid pUCFa (FASMAC Co. Ltd.). We used a cpmApple module with linker sequences (LSS-LI-cpmApple-NH-DQL) from RGECO1, which was a kind gift from Dr. Takeharu Nagai (Zhao *et al*., 2011), for insertion into human DRD1. Sequences coding for hemagglutinin (HA) secretion motif and a FLAG epitope were placed at the 5’ end of the construct as in dLight1.1 (Patriarchi *et al*., 2018) (Fig. 1*A*). *Eco*RI and *Not*I recognition sites were placed at the 5’ and 3’ end, respectively, for subcloning into the expression vector, pCAGGS (Niwa *et al*., 1991) with the ligation by Ligation High ver.2 (TOYOBO). Point mutations of R-GenGAR-DA1, and dLight1.2 and dLight1.3a (Patriarchi *et al*., 2018) were made using polymerase chain reaction (PCR) with the primers containing each mutation and PCR enzyme mixture KOD One (TOYOBO). GRAB_NE1m_ (Feng *et al*., 2019) was provided by Dr. Yulong Li and subcloned into the pCAGGS.

#### Saturation PCR for the screening of optimal linker sequence

To maximize the chromophore fluorescence changes according to the conformational change of R-GenGAR-DA1.0, optimized linker sequences were screened by the saturated PCR. Primers with random bases encoding two-amino acid length were designed as follows. Forward Primer: 5’-TTGCTCAGAAACTTTCAAGTNNBNNBGTGTCCGAAAGAATGTACCC-3’; Reverse Primer: 5’-GTTTCTCTTTTCAACTGATCVNNVNNTGCCTCCCACCCCATAGTTT-3’.

Randomized linker sequences and cpmApple were amplified by PCR and inserted into pUCFa-DRD1-cpmApple plasmid with NEBuilder (NEB). This mutant library was transformed into *E.coli* and the plasmid library was prepared from the mixture of transformed *E.coli*. Library plasmids were digested by *Eco*RI and *Not*I to extract library insert. Library inserts were subcloned into the pCAGGS vector by ligation and transformation into *E.coli*. Single *E.coli* colonies were picked up and the plasmids were prepared from them. Each plasmid was transfected into HeLa cells seeded in 96-well glass-bottom plate with 293-fectin (Thermo Fisher Scientific). Two days after the transfection, cells were imaged as described below.

#### Design of DRD1 mutations based on structural models

Structural models of DRD1 with DA and NE were constructed using PyMOL (The PyMOL Molecular Graphics System, Version 2.0 Schrödinger, LLC) from a crystal structure of the related β_2_-adrenoceptor with bound epinephrine (Ring *et al*., 2013) downloaded from the RCSB Protein Data Bank web site (http://www.pdb.org; PDB code, 4LDO). The binding site residues with side-chain atoms within 5 Å of epinephrine’s aliphatic hydroxy group was exchanged for those of DRD1 by selecting high-probability backbone-dependent rotamers suggested by the mutagenesis wizard in PyMOL. DA was built by deleting the additional methyl plus aliphatic hydroxy group and NE by deleting only the methyl group. Using the same cut-off as above, Ser 107, Val 317 and Trp 321 were identified as residues that could potentially interact with the extra hydroxy group on NE. Asp 103 was disregarded as it is essential for binding of both agonists by interacting with the protonated amine.

With the aim of lowering the binding affinity of NE by removing a potential hydrogen bond to the aliphatic hydroxy of NE, Ser 107 was mutated to Cys and Ala. Additionally, to introduce steric hinderance around the aliphatic hydroxy group, Ile, Leu, Met and Val mutations were also performed. Val 317 was mutated to other hydrophobic residues with longer side chains, that is, Ile, Leu, Phe and Met, again to introduce steric hinderance around the hydroxy in NE. Trp 321 was first mutated to Phe to remove the hydrogen bonding possibility whilst maintaining aromaticity, but since this was detrimental to DA and NE binding, we attempted other residues that maintained hydrogen bonding possibility, that is, His and Gln.

#### Cell culture

HeLa cells were purchased from the Human Science Research Resources Bank. HeLa cells were cultured in DMEM (Wako) supplemented with 10% fetal bovine serum (Sigma-Aldrich) at 37 °C in 5% CO_2_. HeLa cells (3 × 10^4^ cells/well) were plated on CELLview cell culture dishes (glass bottom, 35 mm diameter, 4 compartments; The Greiner Bio-One) (*SI Appendix*, Fig. S4*A*) one day before transfection. Transfection was performed by incubating the cells with a mixture containing 250 ng DNA and 0.25 μl 293fectin transfection reagent (Thermo Fisher Scientific) per well for 4-6 h. Imaging was performed 2 days after transfection.

Primary cultures of rat hippocampal neurons were prepared similarly to that described previously (Fukata *et al*., 2013). Pregnant Wistar/ST rats were purchased from Japan SLC, Inc. A pregnant rat with embryonic rats (embryonic days 19) was killed by CO_2_ inhalation and then embryos (10 embryos per pregnant rat) were removed and decapitated. Hippocampi were dissected from embryonic rat brains and placed in a 10 cm dish on ice with a Hanks’-buffered saline (Ca^2+^/Mg^2+^ free; CMF-HBSS) containing: Hanks’ Balanced Salt solution (Sigma-Aldrich), 10 mM glucose, and 10 mM Hepes (pH 7.4). To dissociate hippocampal neurons, hippocampi were treated with 10 units/ml papain (Worthington Biochemical) for 10 min at 37 °C. Dissociated neurons were plated onto poly-L-lysine (Sigma-Aldrich)-coated 35 mm-glass bottom dishes (3 × 10^5^ cells/well) (*SI Appendix*, Fig. S4*A*) with a plating medium containing: neurobasal medium (ThermoFisher Scientific), 10% FBS, and 10 mM Hepes (pH 7.4). Neurons were incubated at 37 °C and 5% CO_2_ for 3 h, and then the medium was replaced by a medium containing: neurobasal medium, B-27 supplement (ThermoFisher Scientific), 2 mM GlutaMax supplement-I (ThermoFisher Scientific), and 10 mM Hepes (pH 7.4). Half of the medium was removed and replaced with fresh medium every 7 days. The cultured neurons were transfected at 14-21 days *in vitro* by Lipofectamine 2000 (Thermo Fisher Scientific) and were imaged 4-6 days after transfection.

#### Fluorescence imaging

For the imaging of HeLa cells, the medium was changed to imaging buffer [FluoroBrite D-MEM (FB), Life Technologies] supplemented with 1% GlutaMAX (Life Technologies), and 0.2% fetal bovine serum at least 2 h before imaging. For primary hippocampal neurons, the medium was changed to HBSS [119 mM NaCl, 5 mM KCl, 2 mM CaCl_2_, 25 mM Hepes (pH 7.4), 2 mM MgCl_2_, and 33 mM D-glucose] before imaging started.

For the screening of optimal linkers, HeLa cells transfected with library plasmids were imaged with a high content imaging system, IXM-XLS (Molecular Device), equipped with an air objective lens (CFI Plan Fluor 10X, NA = 0.30, WD = 16 mm and CFI Plan Apochromat Lambda 20X, NA = 0.75, WD = 1 mm; Nikon), a Zyla 5.5 sCMOS camera (ANDOR) and a SOLA SE II light source (Lumencor). The excitation and fluorescence filter settings were as follows: excitation filter 562/40 (FF01-562/40-25), dichroic mirror 350-585/601-950 (T) (FF593-Di03-25×36), and emission fluorescence filter 624/40 (FF01-624/40-25) purchased from Semrock. Fluorescence changes before and after application of 10 μM DA were imaged by the IXM-XLS (Molecular Device).

Confocal fluorescence imaging of cells were imaged with an IX83 inverted microscope (Olympus) equipped with a sCMOS camera (Prime, Photometrics), an air objective lens (UPLSAPO 20X, NA = 0.75, WD = 0.6 mm or UPLXAPO 20X, NA = 0.8, WD = 0.6 mm; Olympus), an oil objective lens (UPLSAPO 60X, NA = 1.35, WD = 0.15 mm or UPLXAPO 60X, NA = 1.42, WD = 0.15 mm; Olympus) and a spinning disk confocal unit (CSU-W1, Yokogawa Electric Corporation), illuminated with a laser merge module containing 440 nm, 488 nm, and 561 nm lasers. The excitation laser and fluorescence filter settings were as follows: excitation laser, 440 nm [for cyan fluorecent protein (CFP) and fluorescence resonance energy transfer (FRET) with cyclic adenosine monophosphate (cAMP) biosensor)], 488 nm (for NE1m) and 561 nm (for DA1.2); excitation dichroic mirror, DM445/514/640 (for cAMP biosensor; Yokogawa Electric), DM405/488/561 (for NE1m and DA1.2; Yokogawa Electric); emission filters 465-500 nm (CFP for cAMP biosensor; Yokogawa Electric), 500-550 nm (for NE1m and FRET for cAMP biosensor; Yokogawa Electric), and 580-654 nm (for DA1.2; Yokogawa Electric).

#### Compounds used to test fluorescence response

Stock solutions for the compounds were dissolved in the appropriate vehicle and 0.95 ∼ 1 µl in each 1.5-ml microcentrifuge tube was prepared. Compounds were mixed with 0.5 ml imaging buffer from the well and applied to the same well at each time point during the imaging (*SI Appendix*, Fig. S4*B*). For temperature equilibration of the imaging buffer, 0.5 ml of the imaging buffer was transferred from the well into an empty 1.5-ml microcentrifuge tube and then applied to the buffer in the same well; the procedure repeated 5 times (*SI Appendix*, Fig. S4*C*). The procedure for the compound application in the time-lapse imaging is shown in *SI Appendix*, Fig. S5. The ‘ligand’ dissolved in the appropriate vehicle was applied at the imaging time point shown by the arrow; the ‘vehicle’ was applied at the same time point. The ‘control’ only had light exposure for evaluating the effects of photochromism.

#### Detection of cAMP signaling using cAMP biosensor

The cAMP biosensor ‘CFP-Epac-YFP (yellow fluorescent protein)’, which was developed based on previous work (Ponsioen *et al*., 2004), contains monomeric teal fluorescent protein (mTFP), the human RAPGEF3 (EPAC) gene (corresponding to 149-881 a.a.) obtained from HeLa cells with RT-PCR, and mVenus. The cDNA of cAMP biosensor was inserted into pCX4neo vector (Akagi *et al*., 2003). The plasmid was co-transfected with either DRD1-Tango, which was a gift from Dr. Bryan Roth (Addgene kit # 1000000068) (Kroeze *et al*., 2015), DA1.2, or empty vector. The cells were imaged 2 days after transfection. The level of cAMP was calculated by the ratio of CFP to FRET, followed with normalization by the baseline value before DA application.

#### Quantification of imaging and data analysis

We used Fiji, a distribution of ImageJ (Schindelin *et al*., 2012), for the preparation of quantification and measurement of all imaging files. Principally, for all images, background was subtracted and images were registered by StackReg, a Fiji plugin to correct misregistration, if required. Note that the median filter was used for the time-lapse images of the neuron before registration to remove camera noise preventing registration. Then, regions of interests (ROIs) were selected for the first time point in time-lapse imaging or in the images before the compound application, to surround the whole cell body for HeLa cells and a dendrite near the cell body for hippocampal neurons. Mean pixel intensity in ROIs were measured and these data were further analyzed by Python3 (https://www.python.org). In order to normalize the fluorescence changes with the amount of biosensor expression, ΔF/F_0_ was calculated with the intensity before the compound application as F_0_. The fluorescence change (ΔF/F_0_) image is represented as the pseudocolor intensity-modulated display mode, where color represents the relative ratio value, whilst the brightness of the color represents the fluorescence intensity of the source images. To obtain the EC_50_ and the max ΔF/F_0_, dose-response curves were fitted with Hill function by Python package Scipy1.4 (SciPy.org). Note that the Hill coefficient was fixed as 1 because no cooperative binding was expected.

#### Statistical analysis

All data were presented as mean, with error bars indicating ± SEM if not otherwise specified. Statistical analyses were performed using GraphPad Prism8 (GraphPad Software) and Python 3.0 (Python Software Foundation) with SciPy (SciPy.org) and scikit-posthocs (https://scikit-posthocs.readthedocs.io/) packages. Data were analyzed using Mann-Whitney *U*-test; Student’s *t*-test; one-way ANOVA followed by Dunnett’s or Tukey-Kramer’s post hoc tests as appropriate to correct for multiple comparisons; Friedman test followed by Conover-Iman test with the Bonferroni-Holm correction to correct for multiple comparisons. In Fig. S3 *E* and *F*, normality assumption was judged from Shapiro-Wilk test and Q-Q plot and variances among conditions was supposed to be equal by Bartlett test. All statistical tests were two-tailed. The level of significance was set *P* < 0.05.

### Supplementary Figure legends

**Fig. S1.**
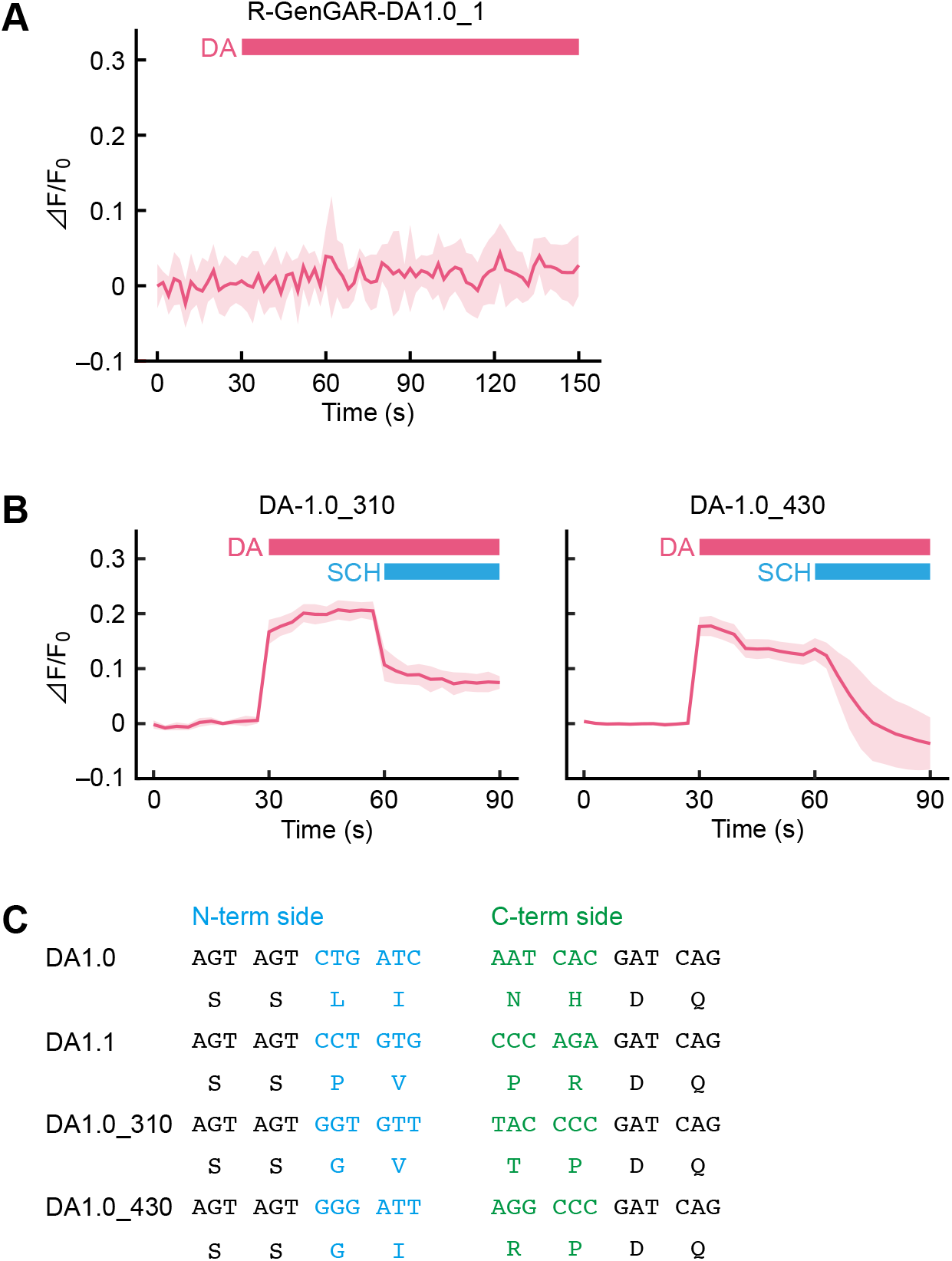
Screening of R-GenGAR-DA1.1. (*A*) Time-lapse imaging of R-GenGAR-DA1.0. Mean ΔF/F_0_ of 10 cells is shown with SD (shaded area). Dopamine (DA, 10 μM) was applied at the time point shown by the pink bar. (*B*) Time-lapse imaging of DA1.0_310 and DA1.0_430. Mean ΔF/F_0_ of 10 cells are shown with SD (shaded area). DA (10 μM) and SCH 23390 (SCH, 10 μM) were treated at the indicated time points shown by pink and blue bars, respectively. (*C*) The amino acid sequence of linker sequences for DA1.0, DA1.1 (DA1.0_76), DA1.0_310, and DA1.0_430, which were obtained from 1st screening.

**Fig. S2.**
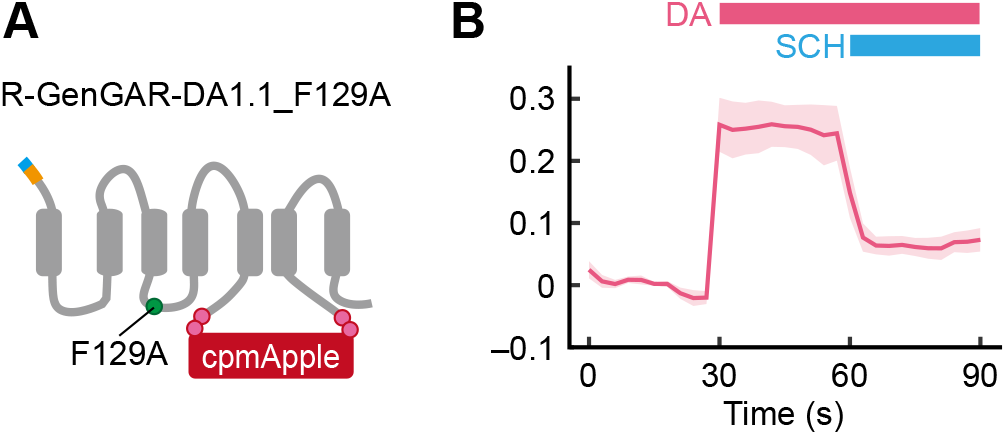
Characterization of R-GenGAR-DA1.1_F129A. (*A*) Schematic illustration of DA1.1_F129A. Phe 129, located in DRD1 intracellular loop 2, mutated to alanin (F129A). (*B*) Time-lapse imaging of DA1.1_F129A. DA (10 μM) and SCH (10 μM) were treated at the indicated time points. Mean ΔF/F_0_ of 20 cells from 2 independent experiments are shown with SD (shaded area).

**Fig. S3.**
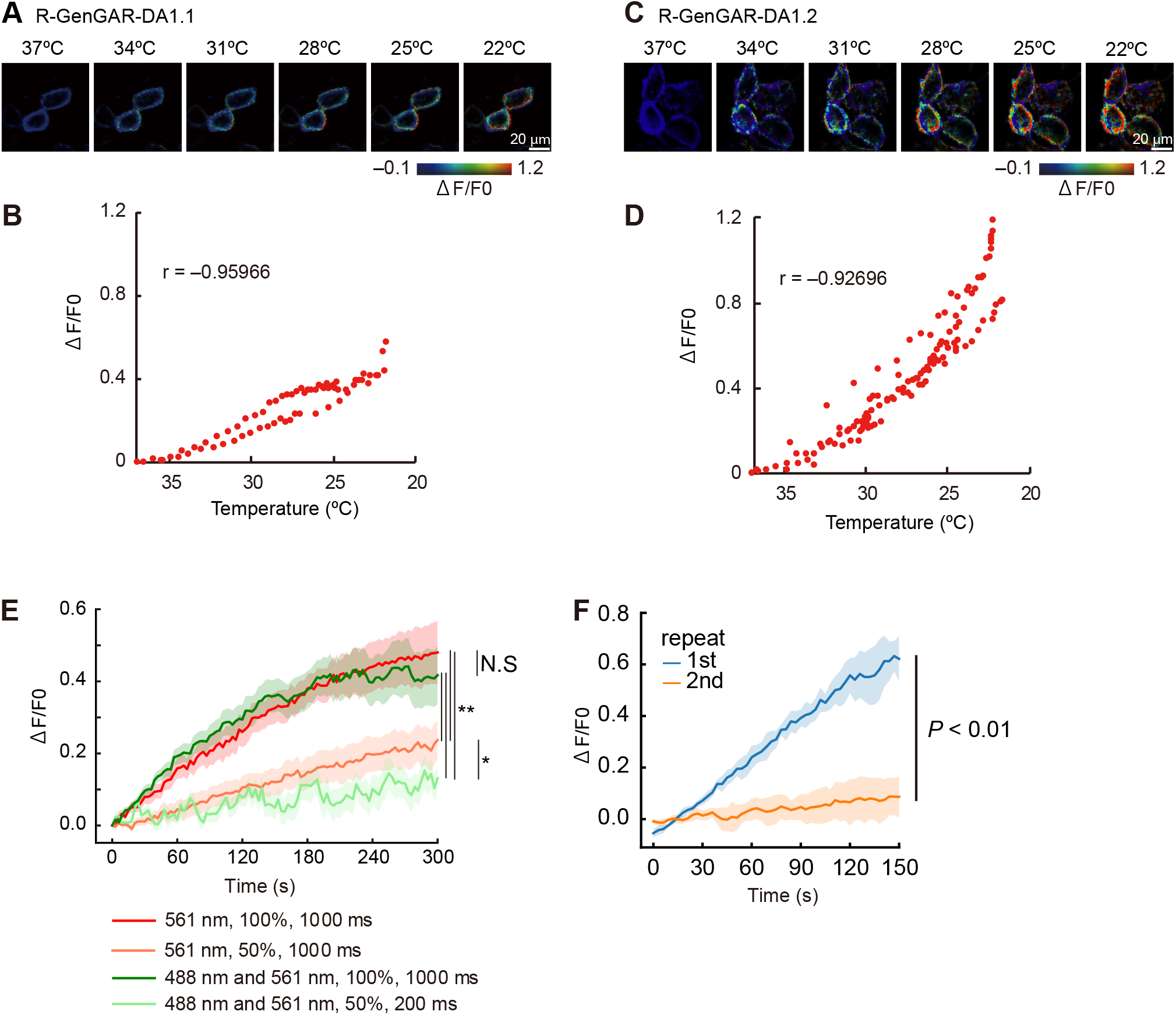
Thermochromism and photochromism for R-GenGAR-DA. (*A–D*) Representative images of HeLa cells expressing DA1.1 (*A*), and DA1.2 (*C*) shown in the pseudocolor intensity-modulated display mode in various incubation temperatures. Regression curve of normalized fluorescence intensity change (ΔF/F_0_) of DA1.1 (*B*), and DA1.2 (*D*) and incubation temperature. Negative correlation between fluorescent intensity and temperature in DA1.1 (r = −0.960, 20 cells in 2 experiments) and DA1.2 (r = −-0.927, 30 cells in 3 experiments) were observed by Pearson product moment correlation coefficient. (*E*) Photochromism-induced change in fluorescence intensity of HeLa cells expressing DA1.2 under the indicated conditions (excitation light wavelength, excitation light power, and exposure time). Incubation temperature was constant during time-lapse imaging. Images were taken every 3 s. The colored-lines represent the average values with the SD of them (shaded area) (*n* = 10 cells in each case). Differences amongst area under the curves (AUCs) from 4 exposure conditions were tested as follows. Normality assumption was judged from Shapiro-Wilk test and Q-Q plot. Variances among conditions was assumed equal following Bartlett test (*P* = 0.696). One-way ANOVA was performed (*F*_3,36_ = 110, *P* < 0.01). As a post-hoc analysis, Tukey-Kramer was used for multiple comparisons (\**P* < 0.05, \*\**P* < 0.01). (*F*) Repeated time-lapse imaging of DA1.2 in primary hippocampal neuron without application of any compounds. Light irradiation protocol as follows: 1-s exposure of 561 nm followed by 1-s exposure of 488 nm; every 3 s for a duration of 150 s. Mean ΔF/F_0_ values of first (blue) and second (orange) 150-s imaging are shown the SD of them (shaded area) (*n* = 4 neurons). Although the mean ΔF/F_0_ values in first 150-s imaging increased gradually because of the photochromism, those in second 150-s imaging were relatively constant and stable. Difference between AUC of first and second imaging was tested by a two-tailed paired *t*-test (*P* = 0.007).

**Fig. S4.**
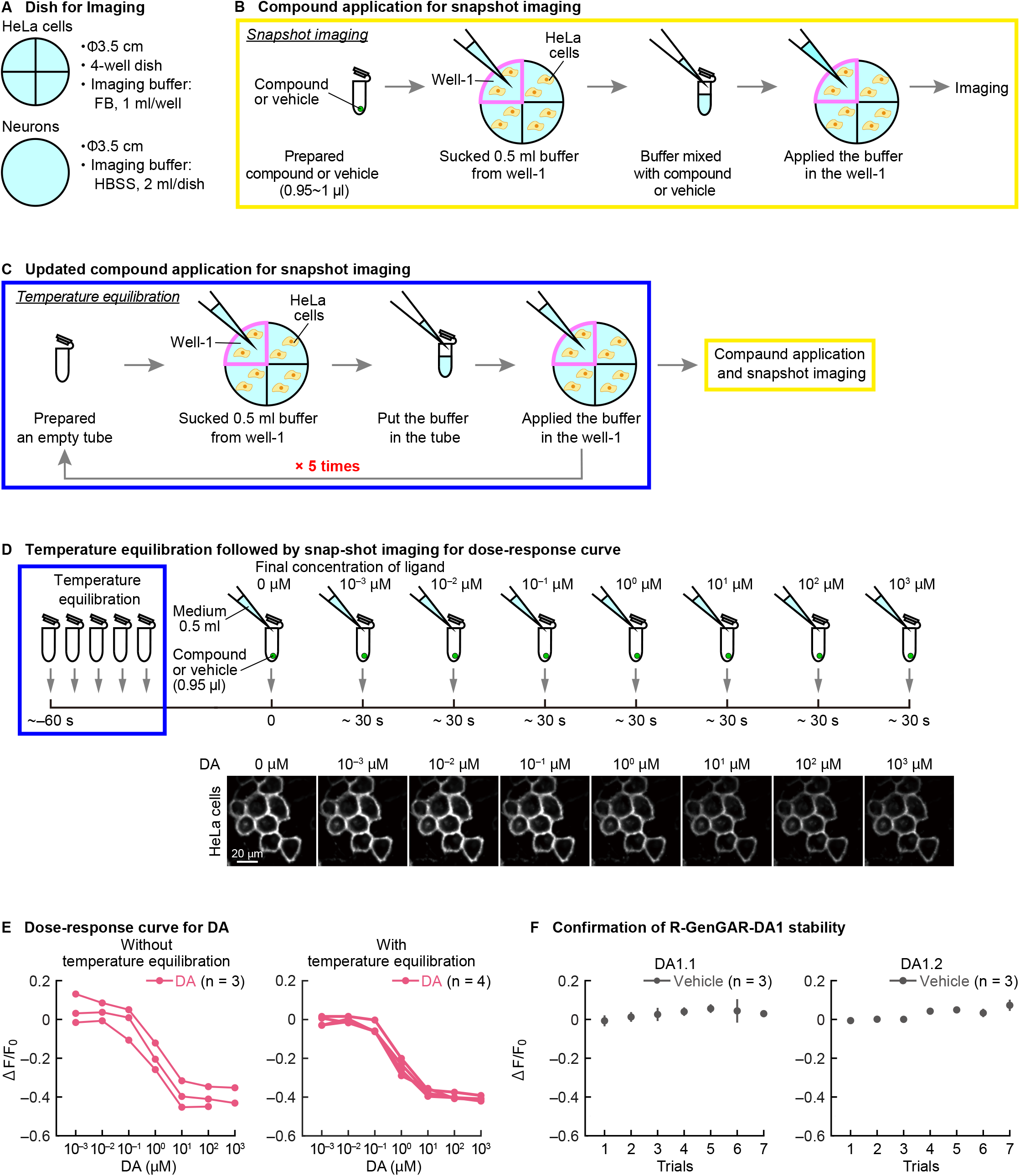
Optimized experimental procedure for the dose-response curve. (*A*) Dishes for imaging of HeLa cells and primary hippocampal neurons. (*B*) Application of compounds for imaging. The compounds, mixed with the 0.5 ml imaging buffer from the well of interest, was applied at the time of imaging. (*C*) Temperature equilibration for imaging. Imaging buffer (0.5 ml out of 1 ml for HeLa cells, and 2 ml for the neurons) from the well of interest transferred to the empty 1.5-ml microcentrifuge tube and returned to the same well; repeated five times. This procedure gradually equilibrated the temperature of the imaging buffer to room temperature and effected the basal fluorescence level of DA1.1 and DA1.2 stable. (*D*) Procedure for making the dose-response curve. Top: the time course of ligand application and imaging shown by the arrow after temperature equilibration. Application of the diluted ligand, imaged sequentially. Bottom: representative images of DA1.2, negatively responding to DA in a dose-dependent manner. (*E*) Quantification of snapshots in the HeLa cells expressing DA1.2 in the dose-response curve for DA without (left) or with (right) temperature equilibration. Temperature equilibration effected to stabilize the basal level ΔF/F_0_ values of DA1.2. (Left, *n* = 4 cells; right, *n* = 3 cells). (*F*) Confirmation of basal stability of R-GenGAR-DA1.1, and DA1.2. HeLa cells expressing DA1.1 (left) or DA1.2 (right) were treated with 7 trials of vehicle stimulation after temperatuire equilibration, showing no change in the mean ΔF/F_0_ values with the SEM of them (*n* = 3 experiments in each). The procedure is the same as *SI Appendix*, Fig. S4*D*.

**Fig. S5.**
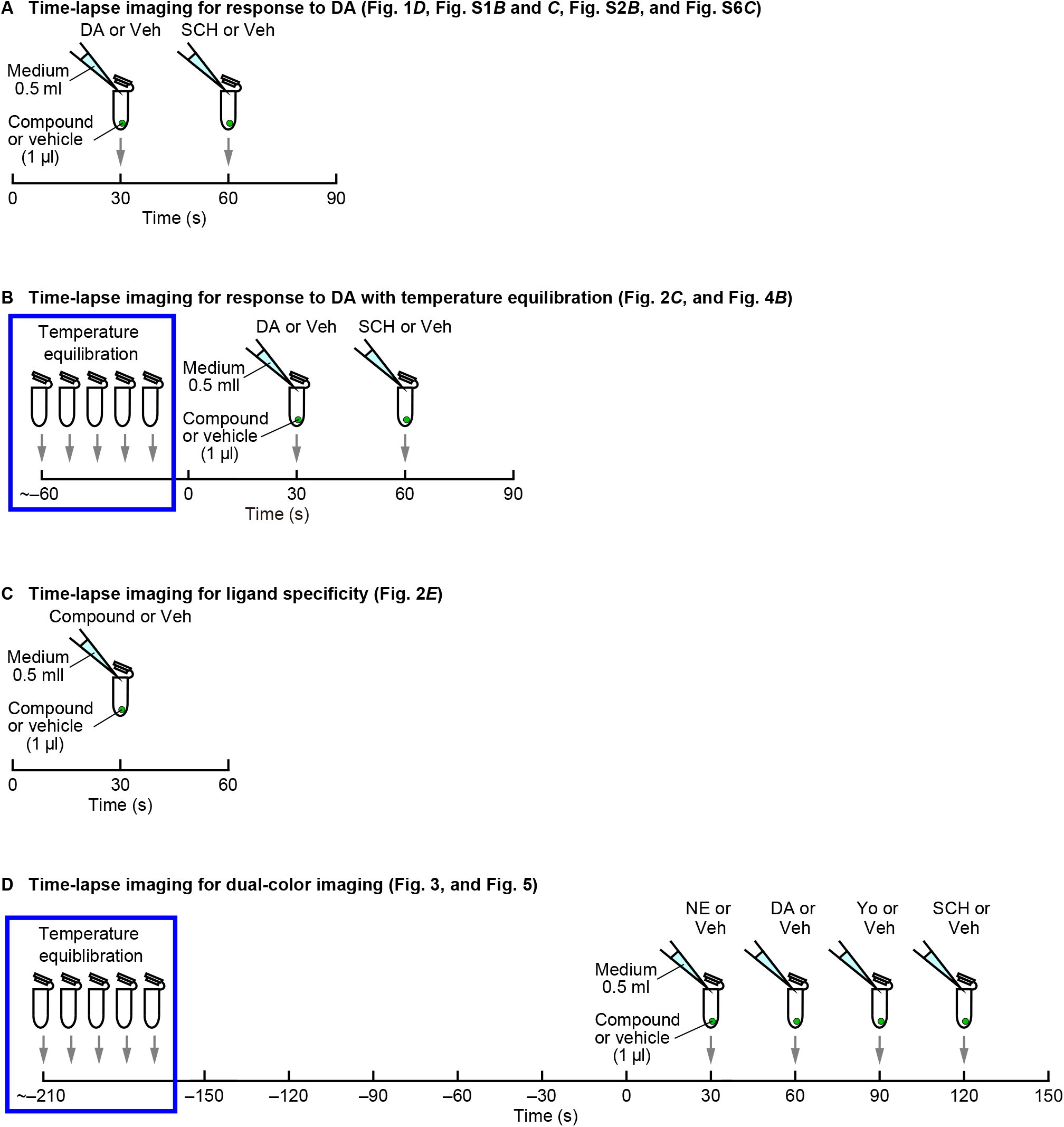
Time course of application of compounds for imaging in HeLa cells and primary hippocampal neurons. (*A*) Compound application for time-lapse imaging without temperature equilibration. Compound or vehicle applied with imaging buffer from the well of interest shown by the arrow. Cells were imaged with the appropriate time exposure (*SI Appendix*, Table S1) acquired every 3 s for a duration of 90 s. (*B*) Compound application with temperature equilibration for time-lapse imaging. Before compound application, temperature equilibration conducted as shown in *SI Appendix*, Fig. S4*C*. Cells were imaged with the appropriate time exposure (*SI Appendix*, Table S1) acquired every 3 s for a duration of 90 s. (*C*) Compound application for checking pharmacological selectivity of DA1.2. Cells were imaged with the appropriate time exposure (*SI Appendix*, Table S1) acquired every 3 s for a duration of 60 s. Averaged ΔF/F_0_ during 30-60 s of each compound was shown in Fig. 2E. (*D*) Compound application for dual-color imaging. After temperature equilibration, we conducted dual-color light irradiation (1-s exposure of 561 nm followed by 1-s exposure of 488 nm light irradiation) every 3 s for a duration of 150 s, which reduced the effect of photochromism, before the start of the imaging. Cells were dual-color imaged (561 nm followed by 488 nm light irradiation) with the appropriate time exposure (*SI Appendix*, Table S1) acquired every 3 s for a duration of 150 s.

**Fig. S6.**
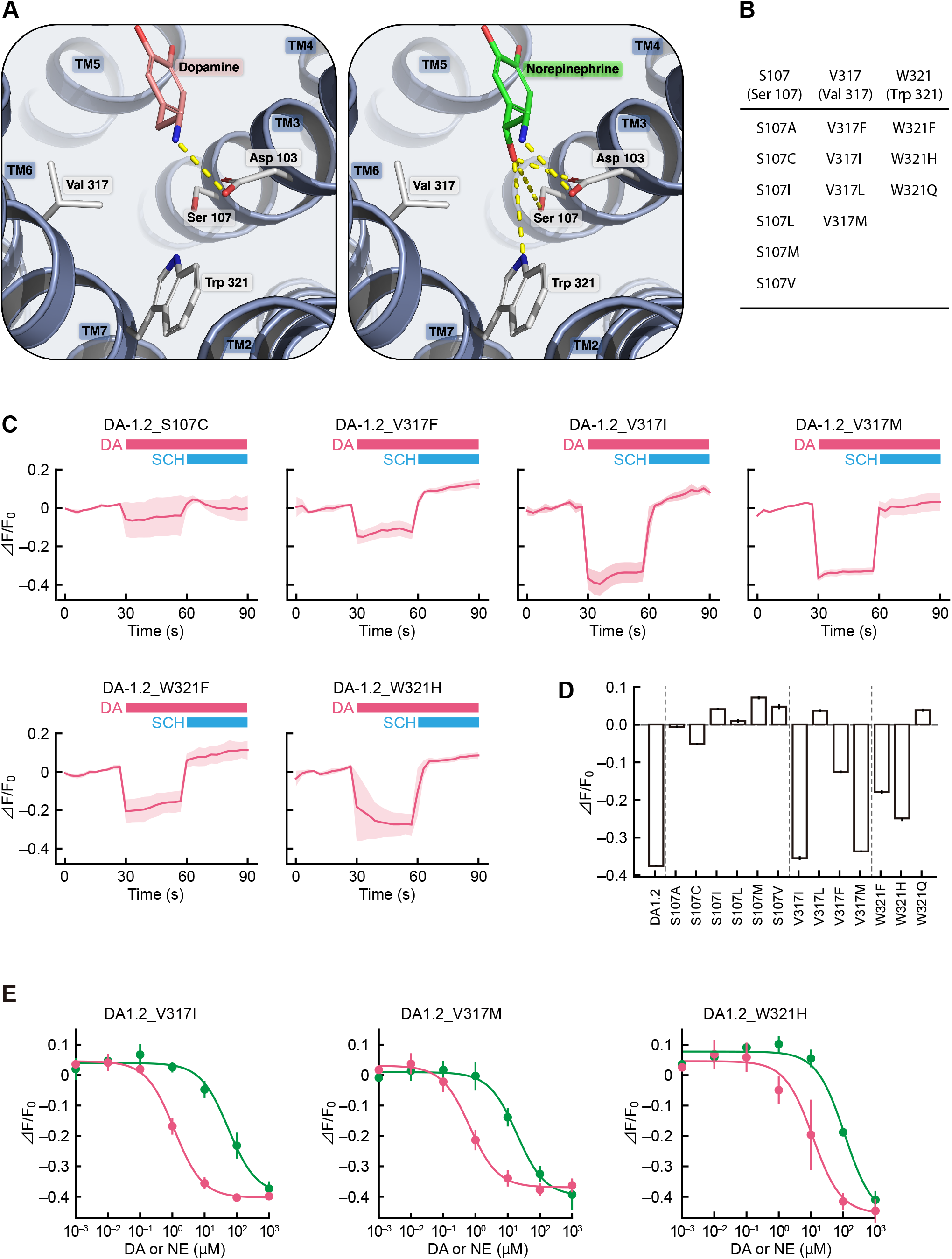
Introducing structural mutations into R-GenGAR-DA1.2. (*A*) Prediction of the residues responsible for the selectivity between DA and NE from structural models of the DRD1 (light blue cartoon and white sticks) with either DA (left, salmon sticks) or NE (right, green sticks) in the binding site. The amino acids close to the additional hydroxy of NE (i.e. Ser 107, Val 317 and Trp 321) may be utilized to affect the preference for binding of DA over NE, e.g. by mutation of hydrogen bonding (yellow dotted lines) amino acids with hydrophobic ones. (*B*) Candidates of structural mutation. (*C*) Mean ΔF/F_0_ (20 cells from 2 experiments in each case) are shown with the SD of them (shaded area). DA (10 μM) and SCH (10 μM) were treated at the indicated time points shown by pink and blue bars, respectively. (*D*) Averaged ΔF/F_0_ during DA application (30-s duration) of each mutant was shown as mean ± SEM. (*E*) The dose-response curves with temperature equilibration of DA (pink) and NE (green) in HeLa cells expressing DA1.2_V317I, DA1.2_V317M, and DA1.2_W321H (*SI Appendix*, Fig. S4*D*). DA1.2_V317I: DA: max ΔF/F_0_ = 0.49 ± 0.01% and EC_50_ = 1.10 ± 0.24 µM; NE: max ΔF/F_0_ = 0.47 ± 0.03% and EC_50_ = 55 ± 14 µM; 50-fold selectivity for DA over NE (DA and NE, *n* = 3 experiments in both cases). DA1.2_V317M: DA: max ΔF/F_0_ = 0.43 ± 0.02% and EC_50_ = 0.66 ± 0.11 µM; NE: max ΔF/F_0_ = 0.42 ± 0.02% and EC_50_ = 19.0 ± 4.1 µM; 28.8-fold selectivity for DA over NE (DA and NE, *n* = 3 experiments in both cases). DA1.2_W321H: DA: max ΔF/F_0_ = 0.55 ± 0.06% and EC_50_ = 12.0 ± 7.4 µM; NE: max ΔF/F_0_ = 0.62 ± 0.07% and EC_50_ = 111 ± 12 µM; 9.3-fold selectivity for DA over NE (DA and NE, *n* = 3 experiments in both cases). Experimental data (dots) were fitted with the Hill equation (lines).

**Fig. S7.**
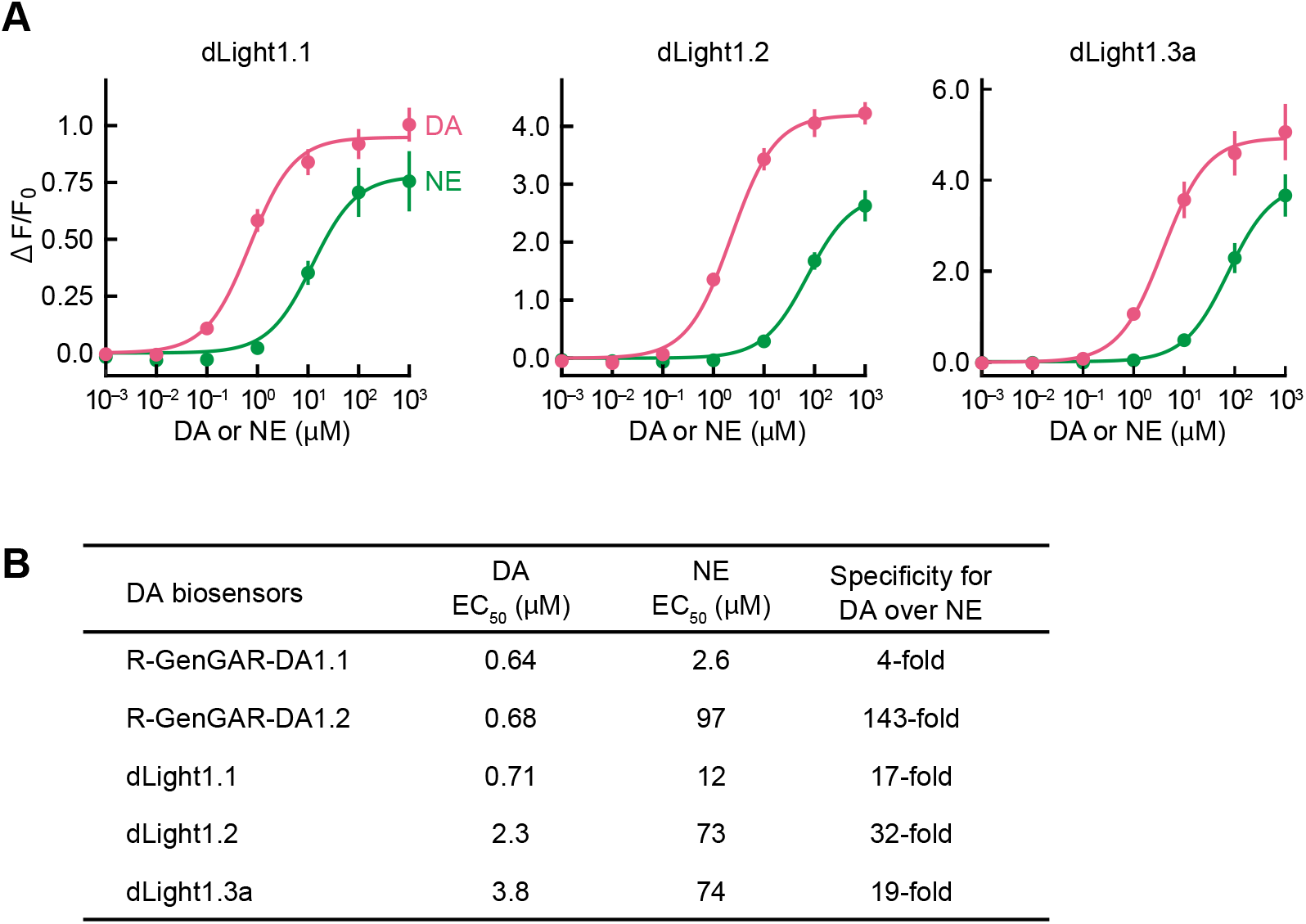
Comparison of selectivity for DA over NE between R-GenGAR-DA and dLight1 sensors. (*A*) Dose-response curve for DA (pink) and NE (green) in HeLa cells expressing dLight1.1, dLight1.2 and dLight1.3a. dLight1.1: DA: max ΔF/F_0_ = 0.95 ± 0.054% and EC_50_ = 0.71 ± 0.083 µM; NE: max ΔF/F_0_ = 0.78 ± 0.11% and EC_50_ = 12 ± 0.55 µM (DA and NE, *n* = 4 independent experiments in both cases). dLight1.2: DA: max ΔF/F_0_ = 4.2 ± 0.19% and EC_50_ = 2.3 ± 0.32 µM; NE: max ΔF/F_0_ = 2.8 ± 0.25% and EC_50_ = 73 ± 5.4 µM (DA and NE, *n* = 4 independent experiments in both cases). dLight1.3a: DA: max ΔF/F_0_ = 4.9 ± 0.50% and EC_50_ = 3.8 ± 0.31 µM; NE: max ΔF/F_0_ = 3.9 ± 0.42% and EC_50_ = 74 ± 4.7 µM (DA and NE, *n* = 4 independent experiments in both cases). (*B*) Summarized affinity for DA and NE, and selectivity for DA over NE of R-GenGAR-DA1.1, R-GenGAR-DA1.2, dLight1.1, dLigh1.2, and dLight1.3a. Selectivity was calculated using EC_50_ of NE relative to EC_50_ of DA.

**Fig. S8.**
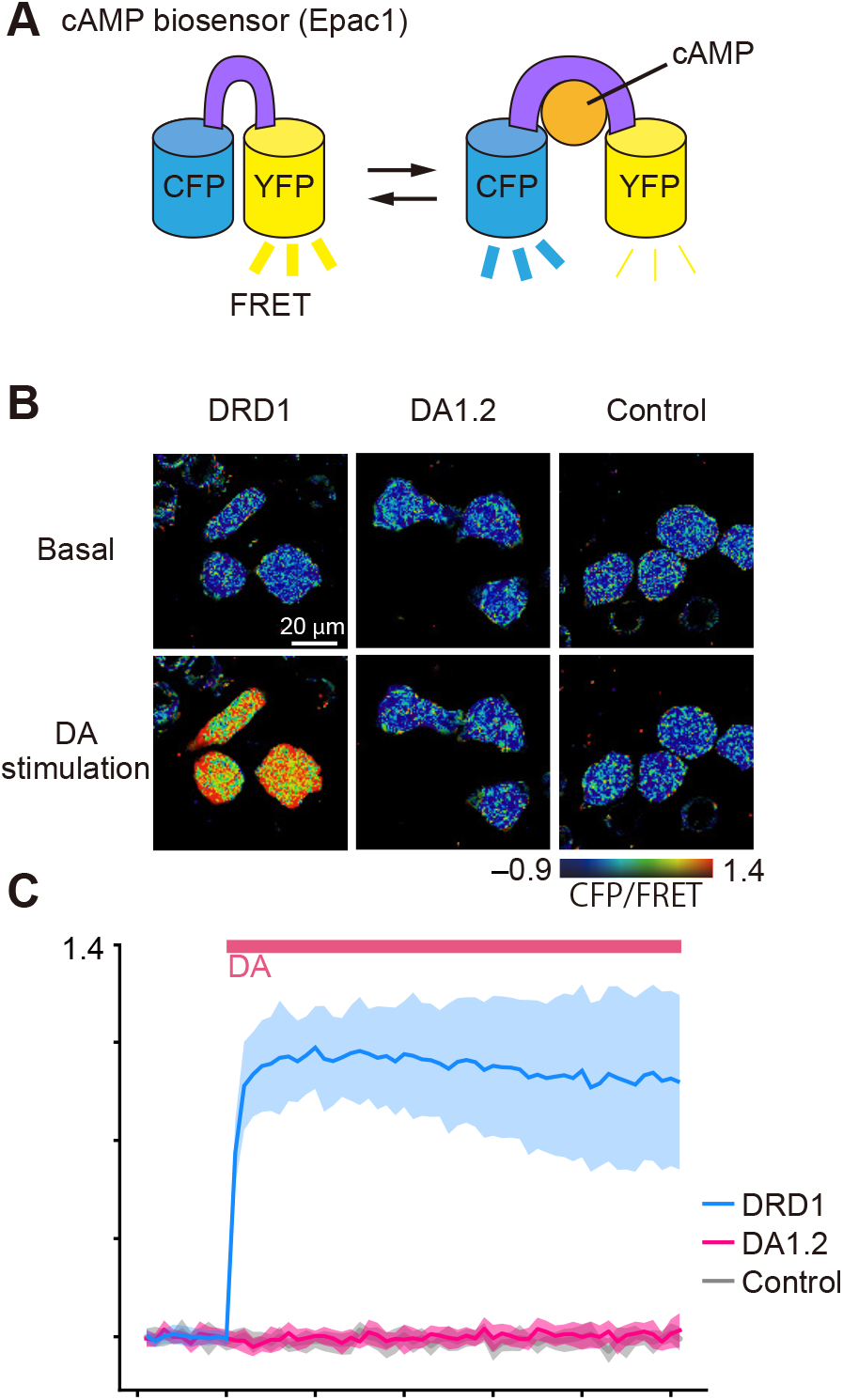
cAMP signaling in HeLa cells expressing R-GenGAR-DA1.2. (*A*) Schematic illustration of the cAMP biosensor, CFP-Epac-YFP. (*B*) Representative images of DRD1 (left), DA1.2 (middle), and control (right, empty vector), which were co-expressing CFP-Epac-YFP, before (top) and after (bottom) application of DA shown in the pseudocolor intensity-modulated display mode. (*C*) Time-lapse imaging of cAMP level (CFP/FRET) in HeLa cells expressing DRD1 (blue), DA1.2 (pink), and control (gray). DA (1 µM) was treated at the time points shown by the pink bar. Cells were imaged with the appropriate time exposure (*SI Appendix*, Table S1) acquired every 1 min for a duration of 30 min. Mean CFP/FRET of 20 cells in 2 experiments is shown with the SD of them (shaded area).

**Fig. S9.**
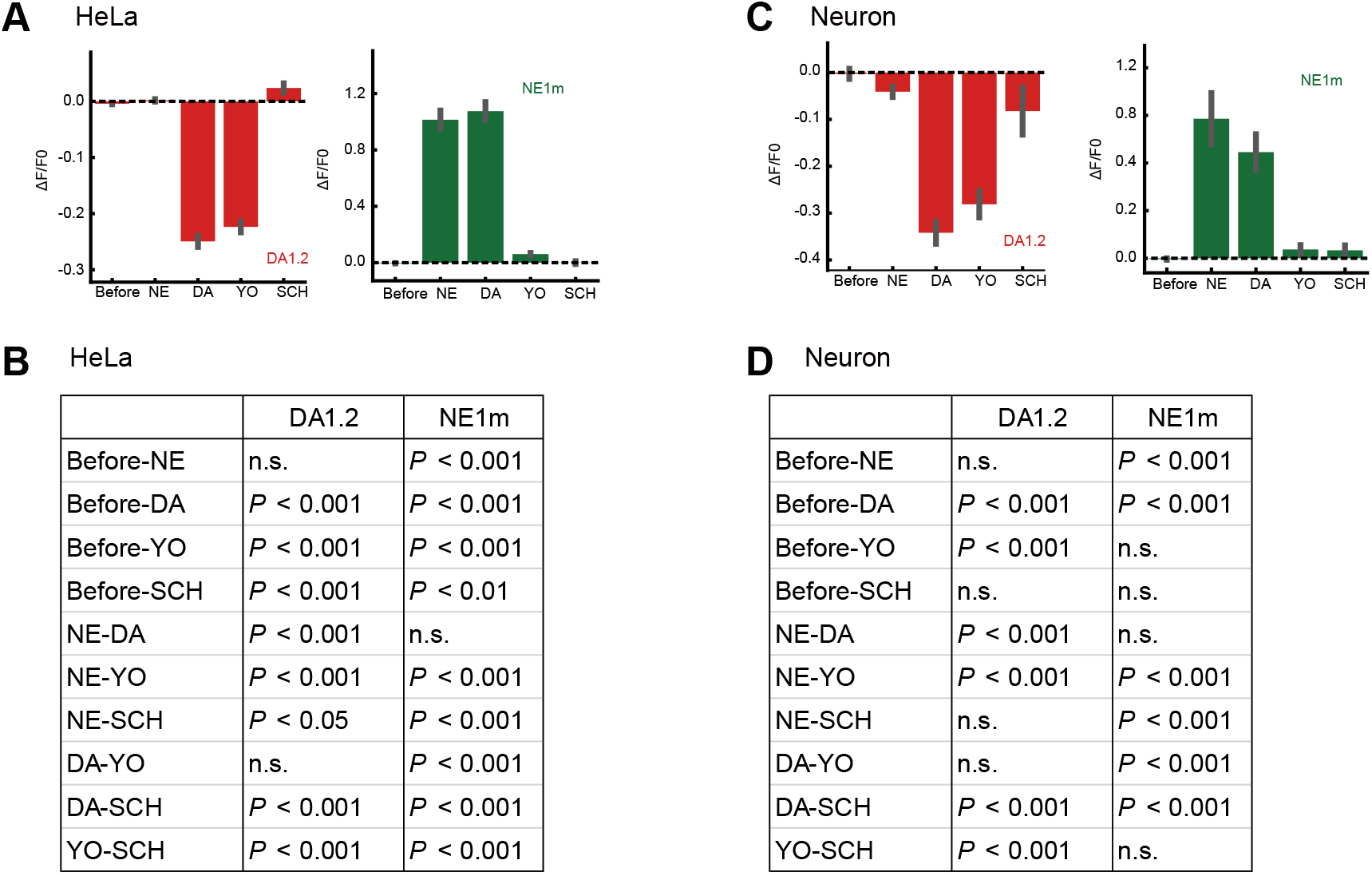
Statistical analysis of dual-color imaging of DA1.2 and NE1m in HeLa cells and primary hippocampal neurons. (*A* and *C*) Quantification of mean values of time-lapse imaging from Fig. 3 (*A*) and Fig. 5 (*C*). Each tiF/F_0_ value for a given compound was normalized by the subtraction of averaged vehicle values along the time course. Bars represent mean ± SEM ΔF/F_0_ values for each consecutive step in the experiment. Each bar represents the mean of the final 15 s (5 time points) of each 30 s condition, which occurs immediately prior to the application of each successive compound. The order of bars from left to right reflects the time course. (HeLa cells *n* = 30 cells, neuron *n* = 6 cells). (*B* and *D*) Statistical results of *SI Appendix* Fig. S 9*A* (*B*) and 9*C* (*D*). There were significant differences between compounds analyzed by Friedman test in HeLa cells (DA1.2, *P* < 0.001; NE1m, *P* < 0.001) and in hippocampal primary neurons (DA1.2, *P* < 0.001; NE1m, *P* < 0.001). Conover-Iman test with the Bonferroni-Holm correction for multiple testing, as a post-hoc analysis, *P* values are shown in the table. n.s., not significant.

**Table S1.** Conditions for fluorescent imaging.

| Figure | Sensor | Cell-type | Microscopy | Filters | Exposure time:<br>laser power |
| --- | --- | --- | --- | --- | --- |
| Fig. 1*B* | DA1.0 | HeLa | IXM-XLS<br>10× (NA = 0.30)<br>20× (NA = 0.75) | Ex: 562/40<br>Dichroic:<br>350-585/601-950 (T)<br>Em: 624/40 | 1000 ms<br>(Lumen cor 100/255) |
| Fig. 1*D*<br>Fig. S1 *A & B*<br>Fig. S2*B* | DA1.0<br>DA1.1 | HeLa | IX83<br>20× (NA = 0.75)<br>20× (NA = 0.80) | Ex: 561 nm<br>Dichroic:<br>DM405/488/561<br>Em: 580–654 nm | 500 ms<br>(Lumen cor 100/255) |
| Fig. 1*E*<br>Fig. 2*B–E*<br>Fig. S3*A–D*<br>Fig. S4*D*<br>Fig. S6*C–E* | DA1.1<br>DA1.2 | HeLa | IX83 with CSU-W1<br>20× (NA = 0.75)<br>20× (NA = 0.80) | Ex: 561 nm<br>Dichroic:<br>DM405/488/561<br>Em: 580–654 nm | 200 ms (ND 100 %) |
| Fig. S3*E* | DA1.2 | HeLa | IX83 with CSU-W1<br>20× (NA = 0.75)<br>20× (NA = 0.80) | Ex1: 561 nm<br>Ex2: 488 nm<br>Dichroic:<br>DM405/488/561<br>Em1: 580–654 nm<br>Em2: 500-550 nm | Ex1: 1000 ms (ND 100%)<br>Ex1: 1000 ms (ND 50%)<br>Ex1, Ex2: 1000 ms (ND 100%)<br>Ex1, Ex2: 200 ms (ND 50%) |
| Fig.3 | DA1.2<br>NE1m | HeLa | IX83 with CSU-W1<br>20× (NA = 0.75)<br>20× (NA = 0.80) | Ex1: 561 nm<br>Ex2: 488 nm<br>Dichroic:<br>DM405/488/561<br>Em1: 580–654 nm<br>Em2: 500-550 nm | Ex1: 200 ms (ND 100%)<br>Ex2: 200 ms (ND 100%) |
| Fig. 4 | DA1.2 | Neuron | IX83 with CSU-W1<br>60× Oil (NA = 1.35)<br>60× Oil (NA = 1.42) | Ex: 561 nm<br>Dichroic:<br>DM405/488/561<br>Em: 580–654 nm | 1000 ms (ND 10 %) |
| Fig. 5<br>Fig. S3*F* | DA1.2<br>NE1m | Neuron | IX83 with CSU-W1<br>60× Oil (NA = 1.35) | Ex1: 561 nm<br>Ex2: 488 nm | Ex1: 1000 ms (ND 10 %)<br>Ex2: 1000 ms (ND 5%) |
|  |  |  | 60× Oil (NA = 1.42) | Dichroic:<br>DM405/488/561<br>Em1: 580–654 nm<br>Em2: 500-550 nm |  |
| Fig. S7 | dLigh1.1<br>dLight1.2<br>dLight1.3a | HeLa | IX83<br>20× (NA = 0.75) | Ex: 488 nm<br>Dichroic:<br>DM405/488/561<br>Em: DM405/488/561 | 500 msec<br>(Lumen cor 20/255) |
| Fig. S8 | CFP-Epac<br>-YFP | HeLa | IX83 with CSU-W1<br>20× (NA = 0.75) | Ex: 440 nm<br>Dichroic:<br>DM445/514/640<br>Em (CFP): 465-500 nm<br>Em (FRET): 500-550 nm | 500 ms (ND 25%) for CFP<br>500 ms (ND 25%) for<br>FRET |

## References

1. J. F. Poulin et al., Mapping projections of molecularly defined dopamine neuron subtypes using intersectional genetic approaches. Nat Neurosci 21, 1260–1271 (2018).

2. S. D. Robertson, N. W. Plummer, J. de Marchena, P. Jensen, Developmental origins of central norepinephrine neuron diversity. Nat Neurosci 16, 1016–1023 (2013).

3. L. A. Schwarz et al., Viral-genetic tracing of the input-output organization of a central noradrenaline circuit. Nature 524, 88–92 (2015).

4. T. Takeuchi et al., Locus coeruleus and dopaminergic consolidation of everyday memory. Nature 537, 357–362 (2016).

5. K. A. Kempadoo, E. V. Mosharov, S. J. Choi, D. Sulzer, E. R. Kandel, Dopamine release from the locus coeruleus to the dorsal hippocampus promotes spatial learning and memory. Proc Natl Acad Sci U S A 113, 14835–14840 (2016).

6. B. S. Beas et al., The locus coeruleus drives disinhibition in the midline thalamus via a dopaminergic mechanism. Nat Neurosci 21, 963–973 (2018).

7. W. Schultz, P. Dayan, P. R. Montague, A neural substrate of prediction and reward. Science 275, 1593–1599 (1997).

8. E. E. Steinberg et al., A causal link between prediction errors, dopamine neurons and learning. Nat Neurosci 16, 966–973 (2013).

9. A. A. Hamid et al., Mesolimbic dopamine signals the value of work. Nat Neurosci 19, 117–126 (2016).

10. A. J. Duszkiewicz, C. G. McNamara, T. Takeuchi, L. Genzel, Novelty and dopaminergic modulation of memory persistence: A tale of two systems. Trends Neurosci 42, 102–114 (2019).

11. B. Panigrahi et al., Dopamine is required for the neural representation and control of movement vigor. Cell 162, 1418–1430 (2015).

12. M. W. Howe, D. A. Dombeck, Rapid signalling in distinct dopaminergic axons during locomotion and reward. Nature 535, 505–510 (2016).

13. B. Xing, Y. C. Li, W. J. Gao, Norepinephrine versus dopamine and their interaction in modulating synaptic function in the prefrontal cortex. Brain Res 1641, 217–233 (2016).

14. Y. Ranjbar-Slamloo, Z. Fazlali, Dopamine and noradrenaline in the brain; Overlapping or dissociate functions? Front Mol Neurosci 12, 334 (2020).

15. S. J. Sara, The locus coeruleus and noradrenergic modulation of cognition. Nat Rev Neurosci 10, 211–223 (2009).

16. M. E. Carter et al., Tuning arousal with optogenetic modulation of locus coeruleus neurons. Nat Neurosci 13, 1526–1533 (2010).

17. A. Eban-Rothschild, G. Rothschild, W. J. Giardino, J. R. Jones, L. de Lecea, VTA dopaminergic neurons regulate ethologically relevant sleep-wake behaviors. Nat Neurosci 19, 1356–1366 (2016).

18. E. Isingrini et al., Resilience to chronic stress is mediated by noradrenergic regulation of dopamine neurons. Nat Neurosci 19, 560–563 (2016).

19. S. Granon et al., Enhanced and impaired attentional performance after infusion of D1 dopaminergic receptor agents into rat prefrontal cortex. J Neurosci 20, 1208–1215 (2000).

20. M. D. Lapiz, D. A. Morilak, Noradrenergic modulation of cognitive function in rat medial prefrontal cortex as measured by attentional set shifting capability. Neuroscience 137, 1039–1049 (2006).

21. M. Wang, S. Vijayraghavan, P. S. Goldman-Rakic, Selective D2 receptor actions on the functional circuitry of working memory. Science 303, 853–856 (2004).

22. S. Vijayraghavan, M. Wang, S. G. Birnbaum, G. V. Williams, A. F. Arnsten, Inverted-U dopamine D1 receptor actions on prefrontal neurons engaged in working memory. Nat Neurosci 10, 376–384 (2007).

23. M. Wang et al., α2A-adrenoceptors strengthen working memory networks by inhibiting cAMP-HCN channel signaling in prefrontal cortex. Cell 129, 397–410 (2007).

24. A. F. Arnsten, S. R. Pliszka, Catecholamine influences on prefrontal cortical function: relevance to treatment of attention deficit/hyperactivity disorder and related disorders. Pharmacol Biochem Behav 99, 211–216 (2011).

25. J. M. Beaulieu, R. R. Gainetdinov, The physiology, signaling, and pharmacology of dopamine receptors. Pharmacol Rev 63, 182–217 (2011).

26. O. Borodovitsyna, M. Flamini, D. Chandler, Νoradrenergic modulation of cognition in health and disease. Neural Plast 2017, 6031478 (2017).

27. P. Devoto, G. Flore, P. Saba, M. Fa, G. L. Gessa, Stimulation of the locus coeruleus elicits noradrenaline and dopamine release in the medial prefrontal and parietal cortex. J Neurochem 92, 368–374 (2005).

28. J. Van Schoors et al., An improved microbore UHPLC method with electrochemical detection for the simultaneous determination of low monoamine levels in *in vivo* brain microdialysis samples. J Pharm Biomed Anal 127, 136–146 (2016).

29. D. L. Robinson, B. J. Venton, M. L. Heien, R. M. Wightman, Detecting subsecond dopamine release with fast-scan cyclic voltammetry *in vivo*. Clin Chem 49, 1763–1773 (2003).

30. A. G. Beyene et al., Imaging striatal dopamine release using a nongenetically encoded near infrared fluorescent catecholamine nanosensor. Sci Adv 5, eaaw3108 (2019).

31. T. Patriarchi et al., Ultrafast neuronal imaging of dopamine dynamics with designed genetically encoded sensors. Science 360, eaat4422 (2018).

32. F. Sun et al., Α genetically encoded fluorescent sensor enables rapid and specific detection of dopamine in flies, fish, and mice. Cell 174, 481–496 e419 (2018).

33. J. Feng et al., Α genetically encoded fluorescent sensor for rapid and specific *in vivo* detection of norepinephrine. Neuron 102, 745–761 e748 (2019).

34. . F. Sun et al., New and improved GRAB fluorescent sensors for monitoring dopaminergic activity *in vivo*. bioRxiv https://doi.org/10.1101/2020.03.28.013722 (2020).

35. Y. Zhao et al., An expanded palette of genetically encoded Ca^2+^ indicators. Science 333, 1888–1891 (2011).

36. Y. Shen, M. D. Wiens, R. E. Campbell, A photochromic and thermochromic fluorescent protein. RSC Adv. 4, 56762–56765 (2014).

37. J. A. Ihalainen, P. Riekkinen, Jr., M. G. Feenstra, Comparison of dopamine and noradrenaline release in mouse prefrontal cortex, striatum and hippocampus using microdialysis. Neurosci Lett 277, 71–74 (1999).

38. P. Devoto, G. Flore, Οn the origin of cortical dopamine: Ιs it a co-transmitter in noradrenergic neurons? Current Neuropharmacology 4, 115–125 (2006).

39. A. V. Leopold, D. M. Shcherbakova, V. V. Verkhusha, Fluorescent biosensors for neurotransmission and neuromodulation: engineering and applications. Front. Cell. Neurosci. 13, 474 (2019).

40. L. Ravotto, L. Duffet, X. Zhou, B. Weber, T. Patriarchi, A bright and colorful future for g-protein coupled receptor sensors. Front. Cell. Neurosci. 14, 67 (2020).

41. C. K. Kim, A. Adhikari, K. Deisseroth, Integration of optogenetics with complementary methodologies in systems neuroscience. Nat Rev Neurosci 18, 222–235 (2017).

42. M. Machacek et al., Coordination of Rho GTPase activities during cell protrusion. Nature 461, 99–103 (2009).

43. S. Mehta et al., Single-fluorophore biosensors for sensitive and multiplexed detection of signalling activities. Nat Cell Biol 20, 1215–1225 (2018).

44. Y. Okubo et al., Visualization of Ca^2+^ filling mechanisms upon synaptic inputs in the endoplasmic reticulum of cerebellar Purkinje cells. J Neurosci 35, 15837–15846 (2015).

45. M. Inoue et al., Rational engineering of XCaMPs, a multicolor GECI suite for *in vivo* imaging of complex brain circuit dynamics. Cell 177, 1346–1360 e1324 (2019).

46. V. Villette et al., Ultrafast two-photon imaging of a high-gain voltage indicator in awake behaving mice. Cell 179, 1590–1608 e1523 (2019).

47. . M. Jing et al., An optimized acetylcholine sensor for monitoring *in vivo* cholinergic activity. bioRxiv https://doi.org/10.1101/861690 (2019).

48. . J. Wan et al., A genetically encoded GRAB sensor for measuring serotonin dynamics *in vivo*. bioRxiv https://doi.org/10.1101/2020.02.24.962282 (2020).

## References

Akagi, T., Sasai, K., and Hanafusa, H. (2003). Refractory nature of normal human diploid fibroblasts with respect to oncogene-mediated transformation. Proc Natl Acad Sci U S A 100, 13567–13572.

Feng, J., Zhang, C., Lischinsky, J.E., Jing, M., Zhou, J., Wang, H., Zhang, Y., Dong, A., Wu, Z., Wu, H., et al. (2019). A genetically encoded fluorescent sensor for rapid and specific in vivo detection of norepinephrine. Neuron 102, 745–761 e748.

Fukata, Y., Dimitrov, A., Boncompain, G., Vielemeyer, O., Perez, F., and Fukata, M. (2013). Local palmitoylation cycles define activity-regulated postsynaptic subdomains. J Cell Biol 202, 145–161.

Kroeze, W.K., Sassano, M.F., Huang, X.P., Lansu, K., McCorvy, J.D., Giguere, P.M., Sciaky, N., and Roth, B.L. (2015). PRESTO-Tango as an open-source resource for interrogation of the druggable human GPCRome. Nat Struct Mol Biol 22, 362–369.

Niwa, H., Yamamura, K., and Miyazaki, J. (1991). Efficient selection for high-expression transfectants with a novel eukaryotic vector. Gene 108, 193–199.

Patriarchi, T., Cho, J.R., Merten, K., Howe, M.W., Marley, A., Xiong, W.H., Folk, R.W., Broussard, G.J., Liang, R., Jang, M.J., et al. (2018). Ultrafast neuronal imaging of dopamine dynamics with designed genetically encoded sensors. Science 360, eaat4422.

Ponsioen, B., Zhao, J., Riedl, J., Zwartkruis, F., van der Krogt, G., Zaccolo, M., Moolenaar, W.H., Bos, J.L., and Jalink, K. (2004). Detecting cAMP-induced Epac activation by fluorescence resonance energy transfer: Epac as a novel cAMP indicator. EMBO Rep 5, 1176–1180.

Ring, A.M., Manglik, A., Kruse, A.C., Enos, M.D., Weis, W.I., Garcia, K.C., and Kobilka, B.K. (2013). Adrenaline-activated structure of β_2_-adrenoceptor stabilized by an engineered nanobody. Nature 502, 575–579.

Schindelin, J., Arganda-Carreras, I., Frise, E., Kaynig, V., Longair, M., Pietzsch, T., Preibisch, S., Rueden, C., Saalfeld, S., Schmid, B., et al. (2012). Fiji: an open-source platform for biological-image analysis. Nat Methods 9, 676–682.

Zhao, Y., Araki, S., Wu, J., Teramoto, T., Chang, Y.F., Nakano, M., Abdelfattah, A.S., Fujiwara, M., Ishihara, T., Nagai, T., et al. (2011). An expanded palette of genetically encoded Ca^2+^ indicators. Science 333, 1888–1891.

